# A single-cell spatial transcriptomic census of human skin anatomy

**DOI:** 10.1101/2025.09.22.676865

**Authors:** Paula Restrepo, Alexis Wilder, Aubrey Houser, Angie Ramirez, M. Grace Hren, Raman Gill, Abiha Kazmi, Larry Chen, Deniz Demircioglu, Dan Hasson, Alan Soto, Stephanie McQuillan, Edgar Gonzalez-Kozlova, Rachel Brody, Benjamin Ungar, Maria Kasper, Phillip Torina, Jesse M. Lewin, Sacha Gnjatic, Sai Ma, Andrew L. Ji

## Abstract

The skin is the largest human organ and a site of significant disease burden, yet its cellular and molecular organization across the body are largely undefined. Here, we construct a spatially-resolved single-cell atlas of 1.2 million cells from normal adult human skin to localize 45 cell types across 15 anatomic sites. We define principles of organ-wide cell composition, including axes of cell diversity and specialization, and distinguish site-enriched cell types. Each body site is comprised of 10 multicellular neighborhoods that define cell-cell communication. Notably, we identify a perivascular neighborhood enriched for immune-stromal crosstalk with features resembling a homeostatic immune niche similar to skin-associated lymphoid tissue. Finally, mapping these neighborhoods onto skin disease reveals pathogenic neighborhood disruptions, including pan-disease immune alterations in the perivascular neighborhood. We present a framework charting the skin’s multiscale spatial organization across a molecular to macroanatomic scale. This work advances our understanding of organ-wide skin cellular organization and communication, and its architectural disruption in disease.

**HIGHLIGHTS:**

- Human skin MERFISH spatial atlas of 1.2 million cells from 114 samples and 22 donors
- Cellular diversity varies along a central to peripheral body site spatial axis
- 10 multicellular neighborhoods define skin microanatomy and homeostatic interactions
- Human skin diseases feature spatial and transcriptional neighborhood remodeling

## INTRODUCTION

Human skin serves critical roles such as thermoregulation and defense against external insults and pathogens. Across the body, different anatomic sites have distinct physiological properties that vary in terms of epidermal thickness, hair follicle density, moisture, pH, temperature, and lipid composition, suggesting functional specialization.^1–3^ Acral surfaces, for example, contain thicker epidermis that provides improved mechanical resistance, but the mechanistic underpinnings of this thickening, along with other phenotypic variation, are not well understood. Diversification of the skin’s cell type composition across the body may facilitate functional adaptation but may also result in unique susceptibilities to chronic disease. Indeed, each body site exhibits well known differences in predilection for skin diseases, which comprise an outsized global burden and negatively impact quality of life, mental health, and socioeconomic status.^4,5^ Atopic dermatitis, for example, often appears on flexural areas, which feature increased pH and reduced water retention, possibly contributing towards a predilection for these sites.^6–8^ Site-specific microbial communities, such as *P. acnes* strains on sebaceous regions like the face, contribute to acne predilection.^2^ Additional external environmental factors such as sun exposure help explain the increased prevalence of UV-induced skin cancers arising on the face and neck,^9,10^ but recent work has shown that anatomic positioning impacts oncogenicity in melanoma, indicating that intrinsic factors can also influence anatomic site-specific disease risk.^11^

Despite these observations, the underlying cellular and molecular programs that maintain the skin’s unique spatial microenvironments are poorly understood. Improving such knowledge could inform broad efforts to improve skin health, from tissue engineering to the development of therapies targeting site-specific cell-cell interactions. Advances in skin organoids grown from human pluripotent stem cells (hPSCs)^12^ hold exciting promise, but a notable lack of immune cells and failure to grow terminal hairs limit their full potential as experimental model systems or transplantable therapies and further signify an absence of key microenvironmental factors. Underscoring this potential, a recent human trial reported successful intradermal injection of volar fibroblasts to modify skin thickness,^13^ paving the way for cell therapies in restoring healthy skin. Therefore, improving our understanding of the skin’s microenvironment within and across anatomic sites has immense potential to aid wide-ranging efforts to improve human skin health. Prior efforts to investigate anatomic site diversity of skin have focused on specific cell types, such as fibroblasts, which exhibit positional memory regulated by HOX transcription factors when cultured *in vitro*.^14,15^ However, recent single cell RNA-seq studies of human skin have revealed the vast heterogeneity of epithelial, immune, and stromal subpopulations within the skin microenvironment, including up to ten subpopulations of fibroblasts in the skin defined by single cell RNA sequencing (scRNA-seq),^16–19^ complicating interpretation of the identity of cultured fibroblasts *in vitro*. More recently, a study incorporating bulk RNA-seq profiling from full thickness skin samples highlighted distinct molecular signatures of the head and neck, trunk and extremities, perinium, and palms and soles,^20^ but whether this reflects unique cell abundances or cell state differences amongst the same population remains unclear. scRNA-seq and spatial profiling have offered further insights into these differences but have thus far featured limited comparisons of 2-3 sites. For example, comparisons of face and trunk skin have revealed enrichment of hair follicle-associated facial fibroblasts that can become cancer-associated fibroblasts in basal cell carcinoma (BCC).^21^ Palm and plantar skin contain unique fibroblast and keratinocyte subpopulations when compared to hip skin.^22^ Finally, while some studies leverage donor-matched profiling of multiple sites, most are limited by an inability to account for natural donor-level variation in comparisons of sites from different donors. Thus, assessing inter-tissue heterogeneity across the body can be exceedingly challenging. Despite advances in characterizing site-specific cell types, a complete *in situ* portrait of the skin’s cellular composition, spatial organization, and intercellular interactions across the human body is lacking.

Here, we combine recent advances in single-cell resolution spatial transcriptomics with a unique study design to construct an organ-wide spatial atlas of normal human skin encompassing 1.2 million cells from 15 anatomic sites and 22 donors. Our cohort included nine donors with multiple sites harvested, including seven for whom we profiled 12 sites, enabling investigation of site differences while accounting for covariates such as age and sex, in addition to inter-individual heterogeneity. We optimized MERFISH^23^ in human skin to identify 45 distinct cell types, matching these annotations to an integrated scRNA-seq atlas from 14 previously published studies. We mapped each cell type’s precise spatial location, differential abundance across sites, and correlated these populations with skin thickness and age. While we identify plantar-enriched keratinocyte and fibroblast subpopulations as strong correlates of skin thickness, papillary fibroblasts also emerged as a universal cell type associated with skin thickness. We further define higher-order organization of cells into 10 multicellular neighborhoods that are universally present across sites but vary in abundance. Notably, we highlight a perivascular neighborhood defined by the enrichment of most immune cell types and a *CCL19*+ perivascular fibroblast subpopulation reminiscent of a homeostatic immune niche. These neighborhoods define cell-cell interactions that we analyzed directly in our MERFISH data and extended to a transcriptome-wide ligand-receptor signaling atlas with imputation from scRNA-seq. Finally, mapping these MERFISH neighborhoods onto publicly available spot-based spatial transcriptomics (Visium) of healthy, tumor, and inflamed skin reveal broad disease-associated cellular and transcriptional remodeling in the immune-enriched perivascular neighborhood, suggestive of a spatially compartmentalized therapeutic target. We further provide these data in an interactive browsable web tool (https://rstudio-connect.hpc.mssm.edu/humanskin-spatialcensus/) to aid the advancement of human skin biology and disease studies. The resulting resource will serve as a basis for future studies seeking to untangle how perturbed microenvironment dynamics and site-specific differences may contribute to skin disease.

## RESULTS

### A single-cell spatial atlas of normal human skin

To map the cellular and spatial diversity of human skin within and across anatomic sites, we sampled clinically and histologically normal human skin from 22 donors (10 males, 12 females) (**Table S1**) with two major collection strategies: first, we sampled up to 12 matched anatomic sites from the same individual donor during autopsies for individuals with a post-mortem index (PMI) of less than 24 hours (n = 99 samples from N = 9 donors, including some technical replicates). Second, to enable the analysis of tissues from live donors and anatomic sites such as the face that could not be obtained from autopsy donors, we also collected surgically discarded skin tissue from panniculectomy (n = 6 samples, N = 6 donors) and tumor-free tissue from Mohs reconstruction (n = 9 samples, N = 7 donors). Donor ages ranged from 25-83 years. In addition to matched H&E stains from adjacent sections, we designed two 500-gene MERFISH panels to profile these samples, composed of a combination of canonical and scRNA-seq derived cell type markers to uniquely identify major subpopulations in the skin, as well as ligand-receptor pairs involved in skin homeostatic cell-cell communication (**Extended Data Figure 1a** and **Table S2**). Overall, 304 (54.1%) genes in the panels correspond to cell type markers, 327 (58.2%) genes overlap with either the CellChatDB (n = 286 genes, 50.9%) or Omnipath (n = 284 genes, 50.5%) ligand-receptor databases, and 434 genes (86.2%) were overlapping between the two panels, enabling integration across panels.^24,25^

To profile gene expression, we extensively optimized sample preparation and clearing conditions for FFPE tissue to enhance RNA detection in the skin (**Methods**), which has high ambient RNAse activity^26^ and extracellular matrix (ECM) content, both of which can significantly compromise efficient transcript detection. Using MERFISH, we profiled 114 unique tissue samples from these 22 donors across 15 anatomic sites to detect a total of 105,046,537 transcripts (**Figure 1a**). We performed several measurements of technical robustness. First, the detected MERFISH counts across two runs of adjacent sections from the same sample correlated nearly perfectly (Spearman correlation r = 0.98), demonstrating high consistency in our sample preparation (**Extended Data Figure 1b**). Second, the detection of 434 overlapping genes between the two panels correlated significantly across runs (r = 0.86; **Extended Data Figure 1c**), demonstrating high precision and suitability for joint integration and analysis. Finally, MERFISH RNA counts correlated highly with completely independent bulk RNA-seq human skin data obtained from GTEx (r = 0.77-0.81; **Extended Data Figure 1d**), confirming accurate gene expression measurements. To examine the impact of RNA quality in our samples with data quality, we correlated the detected transcripts with DV200 scores measured for each sample. Unexpectedly, we found no significant DV200 correlation with detected transcripts, suggesting that tissue DV200 is not necessarily a good predictor of data quality for a MERFISH assay (**Extended Data Figure 1e**). However, in our autopsy samples, we found that the post-mortem interval (PMI), the number of hours from death to sample collection, had a modest negative correlation with transcripts per FOV (r = -0.29; **Extended Data Figure 1f**), suggesting that longer PMIs can adversely impact data quality, likely through RNA degradation. Next, we optimized cell segmentation on DAPI and cell-boundary stains to recover 1,201,886 cells after filtering based on cell size and RNA counts (**Extended Data Figure 1g** and **Methods**). On average, cells contained 62.63 ± 78.5 transcript counts and 27 ± 19.87 genes, in line with previously published MERFISH datasets (**Extended Data Figure 1h**).^27,28^ To assess sources of variance in transcript detection (anatomic site, collection source, panel and donor), we performed pairwise correlations of detected transcript density per sample, demonstrating that most variance could be attributed to gene panel and donor **(Extended Data Figure 1i**).

**Figure 1.**
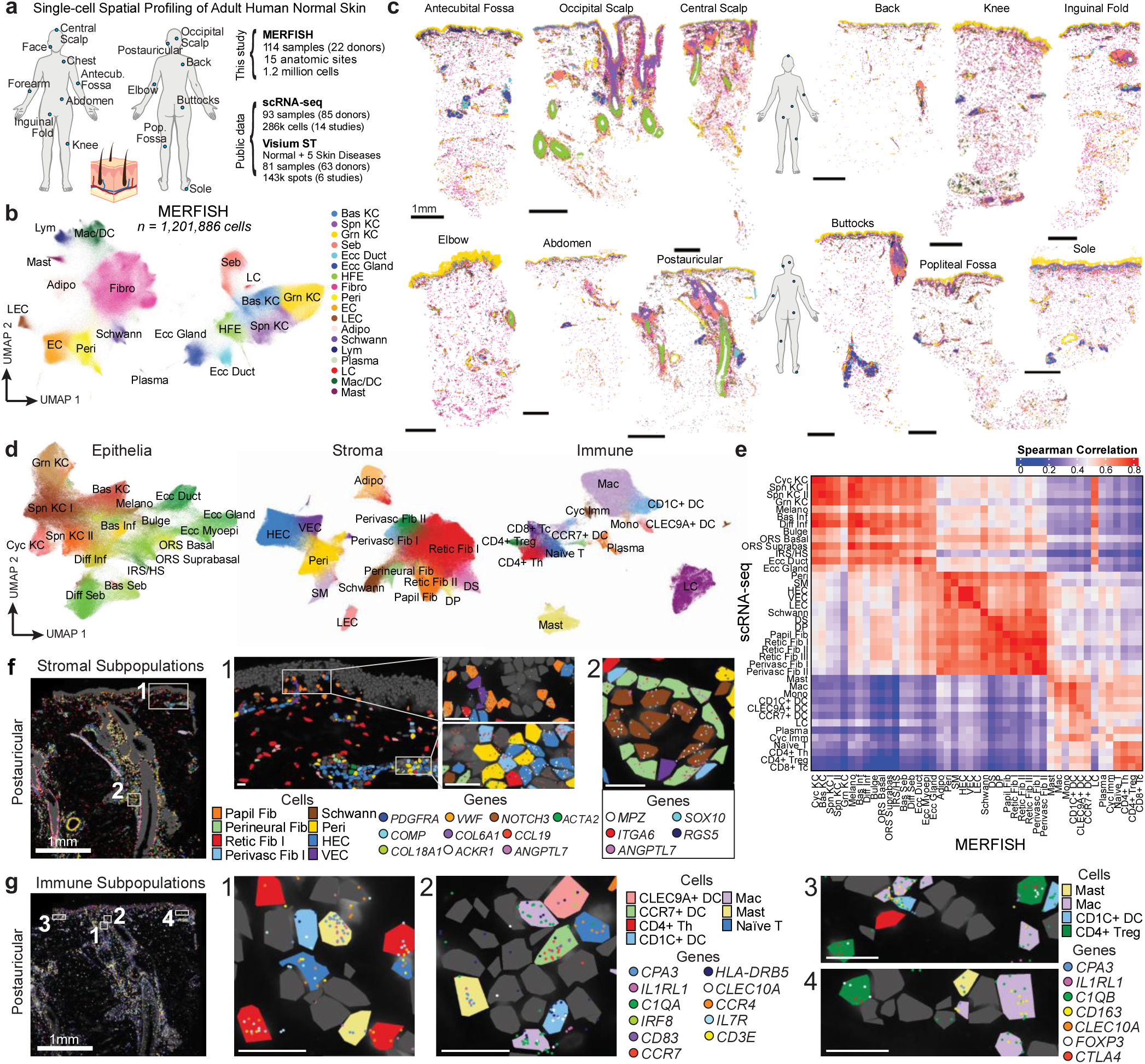
Single-cell spatial transcriptomic profiling of adult human skin using MERFISH. **a.** Overview of the study. **b.** UMAP of integrated MERFISH data from 114 human skin samples labeled by broad cell types. **c.** Single-cell spatial localization of cell types labeled in (B) from 12 anatomic sites from a single donor (D165) profiled with MERFISH. Scale bar = 1mm. **d.** UMAP of epithelial, stromal, and immune subpopulations identified by MERFISH and labeled by cell type. **e.** Pairwise Spearman correlations of overlapping gene expression in MERFISH and scRNA-seq cell types. **f.** Spatial localization of stromal and **g.** immune subpopulations in D165 postauricular sample where each dot represents a transcript.

Integration of our entire MERFISH dataset revealed 18 major cell type populations with distinct spatial localization, and accurate expression of canonical cell type markers (**Figure 1b-c, Extended Data Figure 2a-b, Table S3,** and **Methods**). We observed minimal batch effects from collection source, donor, or gene panel, indicating successful integration (**Extended Data Figure 2c**). To further ensure accuracy of cell type annotations in our MERFISH data, we integrated scRNA-seq data from 14 previously published studies of normal skin samples across 84 donors (n = 285,887 cells; **Extended Data Figure 2d-f** and **Methods**). MERFISH identified broadly similar populations found in scRNA-seq across all studies, (**Extended Data Figure 2a-b** and **2d**), in addition to cell types difficult to dissociate from tissues, such as adipocytes and sebocytes, that were not detected in our scRNA-seq integration (**Extended Data Figure 2a-b** and **Extended Data Figure 2d**).^29^ MERFISH cell type markers correlated well with analogous markers in scRNA-seq data, indicating that our panel accurately identified the same populations in both datasets (**Extended Data Figure 2g** and **Tables S3 and S4**). Finally, the expression of lineage markers in the MERFISH data localized to well-known skin appendages and compartments, such as *SOX9*+ hair-follicle epithelia, *KLF5*+ interfollicular epidermis (IFE) keratinocytes, *COL17A1* and *FLG* localization to the basal and differentiated epidermis, respectively, and various immune and stromal markers in the dermis (*COL1A1*+ fibroblasts, *VWF*+ endothelial cells, *MRC1*+ macrophages) or subcutis (*ADIPOQ+* adipocytes) (**Extended Data Figure 2b**).

To examine data quality differences between living and deceased donors, we compared transcripts per cell and genes per cell across MERFISH runs containing tissues from different collection sources. While deceased donors expectedly exhibited overall lower transcripts and genes per cell (**Extended Data Figure 1h**), likely due to increased RNA degradation, we detected all cell types in autopsy samples, suggesting minimal effects on cell clustering and annotation (**Extended Data Figure 2h**). Together, these data represent the first organ-wide single-cell spatial transcriptomic atlas of adult human skin.

### MERFISH localizes 45 cellular subpopulations in human skin

We next identified and localized cellular subpopulations within our MERFISH data organized into major epithelial, stromal, and immune subpopulations (**Figure 1d**). Expression profiles of all subpopulations identified between both MERFISH and scRNA-seq correlated well, albeit scRNA-seq was notably unable to capture sebocytes, adipocytes, nor eccrine myoepithelial cells (**Figure 1e**). In the epithelial compartment, we identified 17 subpopulations through unbiased clustering, including five IFE keratinocyte (KC) populations corresponding to basal (Bas), spinous (Spn I and II), cycling (Cyc), and granular (Grn) KCs; basal and spinous infundibulum (Bas Inf and Spn Inf); bulge, outer root sheath (ORS) basal and suprabasal; basal and differentiated sebocytes (Bas Seb and Diff Seb); and three cell types in the eccrine (Ecc) compartment, including duct, gland, and myoepithelial (Myoepi) cells (**Figure 1d**). These expressed expected distinct canonical markers (**Extended Data Figure 3a-c**) that also matched analogous clusters in scRNA-seq data (**Extended Data Figure 3a-c**). Visualizing both cell type and marker expression in MERFISH data confirmed the accurate and expected localization of these cell types, such as *COL17A1* and *DST* expression in basal, *DSC1* in spinous, and *FLG* in granular layers (**Extended Data Figure 3a-c**). Consistent with our previous analyses integrating normal skin Visium data with scRNA-seq,^30^ Ecc Duct cells expressed *KRT77* and *GJB2*, while Ecc Gland expressed *DNER* and *SFRP1* (**Extended Data Figure 3c**). Bas Seb was a single layer adjacent to the basement membrane in each sebaceous gland and Diff Seb occupied the suprabasal and central portions of each gland (**Extended Data Figure 3c**).

Notably, we observed two spinous KC clusters, Spn KC I and Spn KC II. Spn KC II was distinguished by elevated expression of *S100A8*, *SOX9*, *GJB2*, and *GJB6* compared to Spn KC I (**Extended Data Figure 3c-g** and **Table S3**). These Spn KC II cells were most prominent in the scalp and sole (**Extended Data Figure 3d**) and exhibited an infundibular-like gene expression profile resembling *S100A8*+ KCs previously described in scalp skin^31^ and *SOX9*+ KCs enriched in palmoplantar skin.^22^ (**Extended Data Figure 3b-c**).^22^ These previously-reported KC subsets have thus far not been compared to one another but were both included in our scRNA-seq reference. Therefore, we next assessed whether these scRNA-seq scalp and sole KCs were transcriptionally distinct. Basal and spinous KCs from both sites clustered together, indicating minimal site-related transcriptional differences, and were clearly distinct from infundibulum clusters, which were enriched in scalp as expected (**Extended Data Figure 3d-g**). Examining basal and spinous KC clusters showed upregulation of *GJB2*, *GJB6*, and *SOX9* in sole compared to scalp, while *S100A8* was more highly expressed in scalp (**Extended Data Figure 3h**). However, it is notable that none of the scalp and sole scRNA-seq data are donor-matched. Thus, while scRNA-seq data can identify differences in gene expression amongst the same cell types across sites, in these data it is: 1) difficult to decouple gene expression differences from donor-level differences and 2) neither scRNA-seq nor MERFISH could readily distinguish scalp- or sole-specific IFE KC subpopulations. As such, we could not confidently sub-cluster Spn KC II further. Taken together, we hypothesize that Spn KC II may represent a uniquely specialized population shared across the two body sites.

We next focused on 15 stromal subpopulations, including fibroblasts, pericytes, endothelial cells (ECs), adipocytes, and Schwann cells (**Figure 1d**, **1f**, and **Extended Data Figure 4a**). Aligning with prior scRNA-seq studies, we identified six fibroblast subpopulations, including papillary (Papil Fib), two reticular (Retic Fib I and II), two perivascular subsets (Perivasc Fib I and II), a newly defined perineural fibroblast subpopulation, in addition to hair follicle associated dermal papilla-like (DP) and dermal sheath (DS) subpopulations (**Figure 1f** and **Extended Data Figure 4a-c**). While prior studies have localized some of these subpopulations, e.g., *CCL19*+ Perivasc Fib I fibroblasts near vasculature,^19^ here we provide definitive localizations across all subpopulations previously observed in scRNA-seq. Notably, *ANGPTL7* marked two of these populations: perineural fibroblasts (*ANGPTL7*+/*ITGA6*+/*KLF5*+) that co-localized with Schwann cells; and Retic Fib II (*ANGPTL7*+/*PRG4*+/*COMP*+) enriched in the sole, consistent with a prior study,^22^ which was scattered throughout the sole dermis **(Extended Data 4c-e)**. Prior studies have identified a fibroblast subpopulation resembling our perineural fibroblasts,^17,22^ and we confirmed expression of these markers (*ANGPTL7*, *ITGA6*, *KLF5*, *MT1X*, *COMP*, *EGFR*) in this previously identified cluster (**Extended Data Figure 4e** and **Table S3**). We also defined two *APOE*+/*CXCL12*+/*C3*+ perivascular populations, distinguished by *CCL19* expression in Perivasc Fib I, which were enriched near superficial blood vessels, while Perivasc Fib II were more enriched around deeper vessels. DP-like cells occupied the epithelial-adjacent regions of the upper hair follicle, while DS cells lined the hair follicles (**Figure 1d**, **1f, and Extended Data Figure 4c**). ECs clustered into lymphatic (LECs, *TFPI*+/*CCL21*+) and two populations of vascular ECs (VECs), one of which was distinguished by *ACKR1* expression, resembling high endothelial venule-like ECs (HECs) (**Figure 1d** and **Extended Data Figure 4a**).^32^

Finally, we identified and localized 13 immune cell types, including DC subtypes (CD1C+, CLEC9A+, and CCR7+ DCs), Langerhans cells (LCs), monocytes, macrophages, mast cells, plasma cells, cycling immune cells, and four T cell subsets (Naïve, CD4+ T helper, CD4+ Tregs, and CD8+ T cells) (**Figure 1d-e** and **Figure 1g**), which also matched scRNA-seq annotations and marker expression (**Extended Data Figure 4f** and **Table S3**). Spatially, while LCs localized in the IFE and hair follicle epithelia (**Figure 1g, Extended Data Figure 3c,** and **Extended Data Figure 4f**), most other immune cells localized to perivascular regions, except for macrophages, which additionally were scattered throughout the dermis (**Figure 1g**). Expression profiles of all 45 subpopulations identified between both MERFISH and scRNA-seq correlated well (notably, scRNA-seq did not capture sebocytes, adipocytes, nor eccrine myoepithelial cells definitively) (**Figure 1g, Extended Data Figure 4f**). Collectively, our MERFISH panels identified and localized 45 unique cell populations in adult skin at a comparable granularity to scRNA-seq, yielding an unparalleled high-resolution single-cell spatial atlas of human skin (**Extended Data Figure 5**).

### Stereotypic patterns of skin cell type diversity, density, and abundances across anatomic sites

Given inter-tissue heterogeneity and functional specialization of skin at distinct sites, we next defined metrics to globally compare tissues across anatomic sites. We therefore quantified the cellular diversity (Shannon diversity index), which serves as a proxy for specialization, and the cell density (cells per 100 µm^2^) of each site. These analyses revealed the highest diversity at hair-dense scalp, face, and postauricular sites, while elbow and knee featured the lowest diversity (**Figure 2a-b**). While higher diversity may reflect an increase in hair follicle-associated populations at face and scalp, interestingly, the sole scored the next highest in diversity despite being a hairless site. Further, while elbow and knee were lowest in diversity, flexural sites of the limbs such as antecubital and popliteal fossae featured higher diversity, highlighting differences despite similar regional coordinates (i.e., extremities). Finally, centrally located body sites such as the buttocks, abdomen, and back were low in diversity, suggesting diversity increases centrifugally from central to peripheral sites. Similarly, density grossly correlated with diversity, with abdomen, back, and buttocks exhibiting the lowest density while face, scalp, and sole had the highest density, suggesting a centrifugal density axis across the body plan (**Figure 2a-b**).

**Figure 2.**
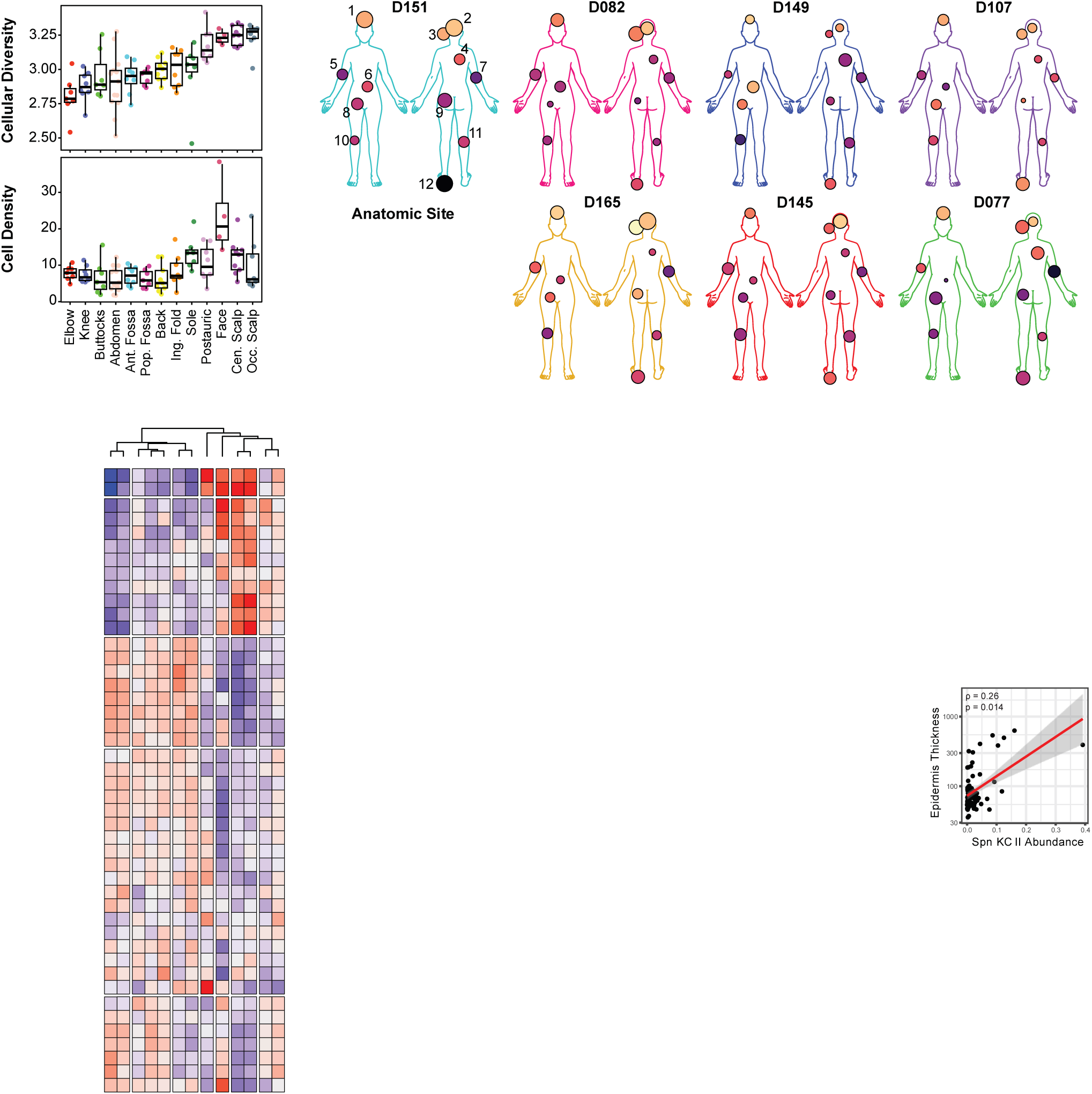
Stereotypic patterns of cell type diversity, density, and abundance across sites. **a.** Box plots of cellular diversity (Shannon diversity index) and density (cells/mm^2^) at each anatomic site, ranked from lowest to highest diversity. Boxes display median and top and bottom quartiles of the data. **b.** Body maps depicting diversity and density for seven donors with 12 anatomic sites profiled. **c.** Left, clustered heatmap of cellular abundances across anatomic sites. Right, bar plots of variance portioning showing contributions of each metadata to the variance in data. -adjusted p < 0.1, *adjusted p < 0.05, **adjusted p < 0.01, ***adjusted p < 0.001, moderated t test. **d.** Box plots of cellular diversity (Shannon diversity index) and density (cells/100 μm^2^) at each anatomic site by tissue compartment, ranked from lowest to highest overall diversity, as in (a). Boxes display median and top and bottom quartiles of the data. **e.** Principal components analysis of tissue compartment density and diversity. **f.** Spearman correlations of cell type proportion and epidermal/stratum corneum thickness. *p < 0.05. **g.** Scatter plots and Spearman correlations of epidermal thickness with Spn KC II, Retic Fib II, and Papil Fib abundances. Each point is a tissue sample (N = 89 samples from 7 donors).

To assess underlying cellular constituents defining diversity, we compared specific cell type abundances across sites using crumblr (**Methods**).^33–35^ Crumblr identified specific cell type enrichment and depletion across sites, clustering both cell types and sites based on cellular composition (**Figure 2c**). Remarkably, clustering of anatomic sites revealed similar patterns of cell type composition within flexural (antecubital and popliteal fossae), extensor (elbow and knee), trunk (back, abdomen, and buttocks), or scalp (central and occipital scalp) sites, while face and sole clustered uniquely by themselves (**Figure 2c**), highlighting the robustness of our study design and donor-matched sampling strategy. High-diversity sites such as the face and scalp featured a wider variance in cell type abundances compared to other sites. Consistent with low diversity patterns, the buttocks, abdomen, and back exhibited minimal enrichment or depletion of any cell type (**Figure 2c**). Collectively, these data suggest distinct but highly stereotypic cell compositions across sites.

Combining these analyses with variance partitioning of covariates such as age, gender, donor, batch, and tissue area imaged of epidermal, dermal, and subcutis compartments confirmed that the cell type abundances largely varied by anatomic site, while quantifying other sources of variance (**Figure 2c**). We further visualized each tissue sample in a UMAP using each cell type abundance as the features, which showed similar clustering by these body categories and trajectories of density and diversity (**Extended Data Figure 6a-c**). Visualizing adjusted centered log-ratios (CLRs) for each cell type across sites demonstrated consistent trends at the donor level (**Extended Data Figure 6d**). Cell types driving anatomic site differences included, expectedly, Spn KC II (scalp and sole enrichment), Retic Fib II (sole), and hair follicle- and sebaceous gland-associated subpopulations (face, scalp, and postauricular) (**Extended Data 3c** and **Figure 2c**). Interestingly, we observed that innate immune cells such as monocytes, macrophages, and DC subsets were strongly enriched in the extremities, with similar trends for vasculature-associated cell types such as VECs, HECs, pericytes, and Perivasc Fib I/II subpopulations, in addition to LECs and Retic Fib I (**Figure 2c**). However, T cell subsets were distinctly enriched in antecubital fossa and to a lesser extent the popliteal fossa, divergent from elbow and knee patterns (**Figure 2c**). In contrast, cell types of the epidermis and eccrine gland were remarkably stable across sites. Together, these analyses reveal how distinct stable and dynamic cell types drive anatomic site differences by forming unique coalitions of cellular components across the skin.

Next, to begin assessing intra-tissue heterogeneity, we manually annotated our tissues by classic histology compartments of the epidermis, dermis, and subcutis, cross-referencing these with matched histology from our cohort, and assigned each cell to its compartment (**Extended Data Figure 6e**). Re-calculating compartmental diversity and density revealed that the dermis is the most overall diverse compartment, largely matching overall tissue diversity patterns, followed by the subcutis and epidermis (**Figure 2d**). Interestingly, the sole’s dermal diversity was among the lowest across sites despite high overall density (**Figure 2a**). Instead, high epidermal and subcutis diversity drove the overall diversity of the sole, suggesting site differences can be driven by distinct compartments. In terms of compartment density, the epidermis was the overall densest, followed by dermis and subcutis, with the face exhibiting both the highest epidermal and dermal density (**Figure 2d**). Principal components analysis of the cell type abundances per compartment showed clustering of samples by compartment, revealing density and diversity as major sources of PC1 and PC2 variance, respectively (**Figure 2e**). Thus, the skin possesses several means to achieve functional specialization, including diversification of distinct compartments that contribute to overall tissue characteristics.

Finally, to correlate cell type abundances with functional tissue characteristics, we leveraged our histology data to measure the average epidermal (basal through stratum corneum) and stratum corneum thicknesses across all our sites (**Extended Data Figure 6f**) and performed Spearman correlation with each cell type proportion in each sample, along with pairwise cell type proportion correlations (**Figure 2f**). Cell type proportion correlations clustered into expected compartments; for example, subpopulations of the IFE, pilosebaceous unit, eccrine glands exhibited high correlations (**Figure 2f**). Regarding epidermal and stratum corneum thicknesses, since the sole is thickest in both regards, sole-enriched cell types such as Spn KC II and Retic Fib II were expectedly significantly correlated with both epidermal and stratum corneum thickness (**Figure 2f-g**). Interestingly, Papil Fib was the only other cell type that exhibited significant correlation with both (**Figure 2f-g**) and was driven by the sole, elbow, and knee (**Figure 2c**), sites that may experience more mechanical stress than others. Beyond the known signaling between Papil Fib and epidermal KCs, this finding was particularly intriguing given a recent report of a successful first-in-human trial of intradermal injections of sole fibroblasts to increase non-sole skin thickness.^13^ Our data suggest that purifying and injecting papillary fibroblasts from sites beyond the sole could potentially also facilitate skin thickening.

### Defining multicellular neighborhoods in human skin

Given stereotypic cell compositions across sites, we sought to determine whether spatial organization also exhibited recurrent patterns in cell type proximity. Spatial proximity enrichment analysis identified clusters of proximal cell types resembling the cell type proportion correlations (**Figure 2f** and **Extended Figure 7a**). For example, Schwann cells and perineural fibroblasts we tightly linked as expected, as were hair follicle bulge, ORS basal/suprabasal, and IRS/HS cells. To further systemically examine intra-tissue heterogeneity in an orthogonal unbiased manner, we performed spatial clustering of our entire dataset using CellCharter.^36^ Iterative clustering showed maximum solution stability at *k* = 10, revealing 10 multicellular neighborhoods present across all samples (**Figures 3a-b, Extended Figure 7b-c** and **Methods**). We annotated these neighborhoods based on visualizing their locations within each sample: N0 was located at the dermo-epidermal junction (DEJ); N1 encompassed perivascular areas both superficial and adjacent to hair follicles, which we named PERIVASC I; N2 was strongly associated with the IFE and contained the differentiated layers (DIFF IFE); N3 also favored perivascular areas but surrounding deeper and larger vessels (PERIVASC II); N4 encompassed the large areas of reticular dermis between vessels (STROMA); N5 strongly associated to the upper hair follicle (UPPER HF); N6 was located at the eccrine glands (ECCRINE); N7 was located at sebaceous glands (SEB GLAND); N8 was predominantly located in the subcutis (SUBCUTIS); and N9 covered the lower hair follicle (LOWER HF) (**Figure 3b** and **Extended Figure 7d**).

**Figure 3.**
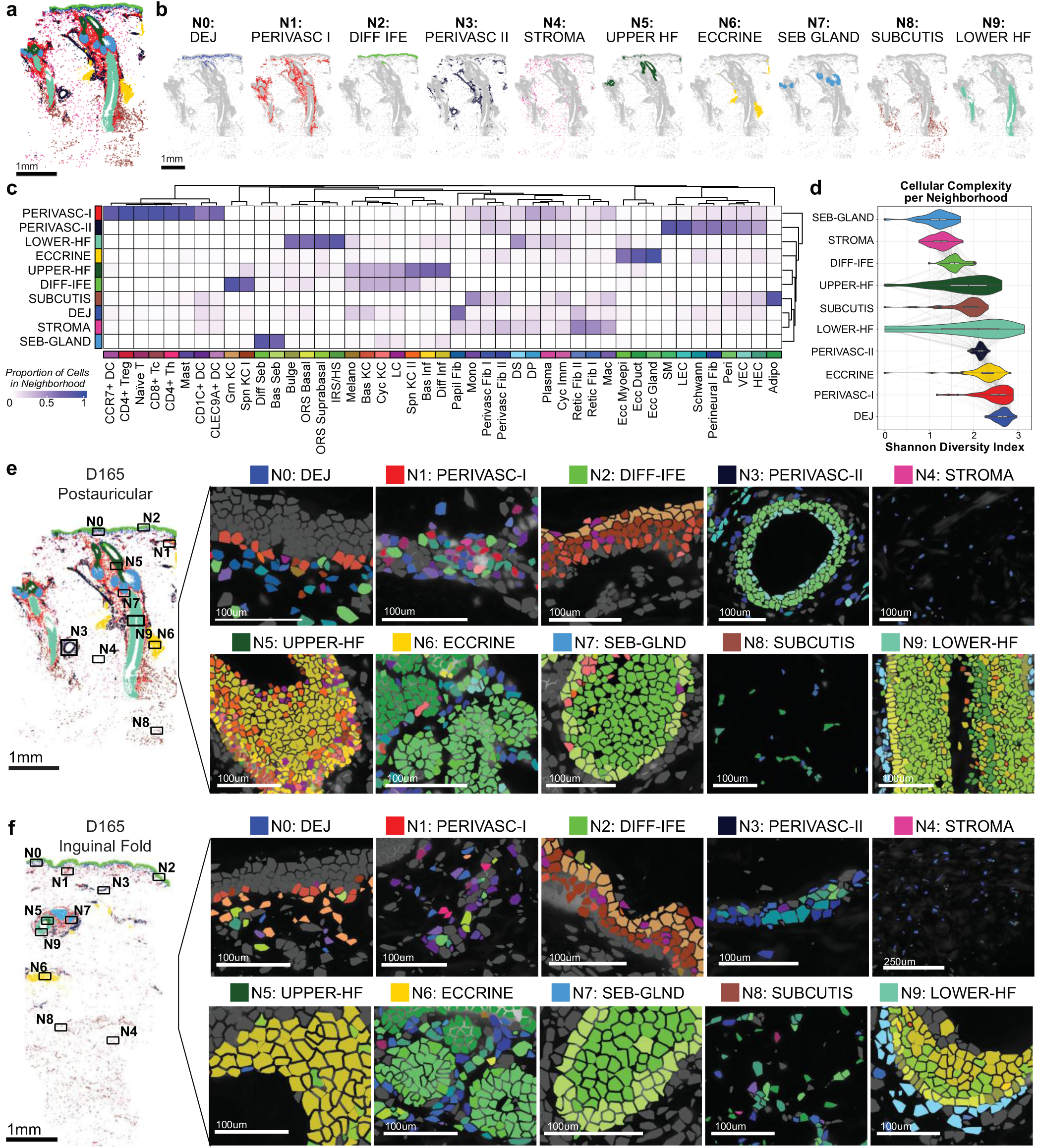
Multicellular spatial neighborhoods define cell composition and localization. **a.** D165 postauricular sample labeled by 10 multicellular neighborhoods. **b.** Spatial highlights of cells labeled by each neighborhood from (A). DEJ = dermo-epidermal junction, DIFF = differentiating, IFE = interfollicular epidermis, HF = hair follicle, SEB = sebaceous. **c.** Clustered heatmap of cell type proportions across neighborhoods. Each column sums to 1. **d.** Violin plots of Shannon diversity indices in each neighborhood (N = 114 samples each). **e.** D165 postauricular neighborhood zoom-ins for cell type constituents of each neighborhood. Cell types are colored by same colors as in (c). **f.** Same as (e) but for D165 inguinal fold sample.

We also examined how distinct cell types were distributed across these neighborhoods and found strong localization preferences for many cell types towards distinct neighborhoods (**Figure 3c**). Expectedly, the canonical cell types of the epidermis, hair follicles, and sebaceous/eccrine glands were enriched in their respective neighborhoods. Interestingly, most immune cells, including DC subsets, T cells, and mast cells, exhibited strong enrichment for the PERIVASC I neighborhood, in addition to Perivasc Fib I/II and vasculature-associated cells. PERIVASC II was composed of vasculature-related cells as well as lymphatic ECs and fewer immune cells. Neighborhoods exhibited a range of diversity scores, with SEB GLAND (composed of mostly Bas and Diff Seb cells) and STROMA (Retic Fib I/II and macrophages) at the low end, and the PERIVASC I and DEJ neighborhoods exhibiting the highest diversity (**Figure 3d**). We visualized the cell types within each neighborhood, confirming their spatial positions and proximity (**Figure 3e-f**). Together, these data define the microanatomy of human skin into recurrent spatial multicellular neighborhoods, suggesting higher order organization of the distinct cell types across the human body.

We next assessed how neighborhood abundance varies across sites. We expected less variation of neighborhoods across sites given that they group cells into fewer components or dimensions than cell types (10 neighborhoods compared to 45 cell types). Nonetheless, neighborhood abundances also recapitulated major body coordinates including clustering of hair-dense scalp/postauricular/face sites, extremities, and trunk, while sole remained unique (**Figure 4a-b**). This variation was driven by a few neighborhoods that exhibited significant enrichment at specific sites, including enrichment of the three pilosebaceous neighborhoods (UPPER HF, SEB GLAND, LOWER HF) in the scalp, while STROMA, DEJ, DIFF IFE, and PERIVASC II were enriched in elbow/knee. Variance partitioning revealed that PERIVASC I and SUBCUTIS were two neighborhoods which remained relatively stable across sites with low percentage of compositional variance explained by anatomic site (**Figure 4b**). Interestingly, the UPPER HF neighborhood was also enriched in sole, which highlighted the transcriptional similarity of sole-enriched Spn KC II cells with Diff Inf cells. Given shared expression of the hair follicle master regulator *SOX9* between these populations in prior studies^22^ and our own data (**Extended Data Figure 3a-c**), these observations may suggest that the expression of *SOX9* or other hair follicle regulators in sole KCs may serve critical functions for this site’s increased thickness and unique ability to handle mechanical load.

**Figure 4.**
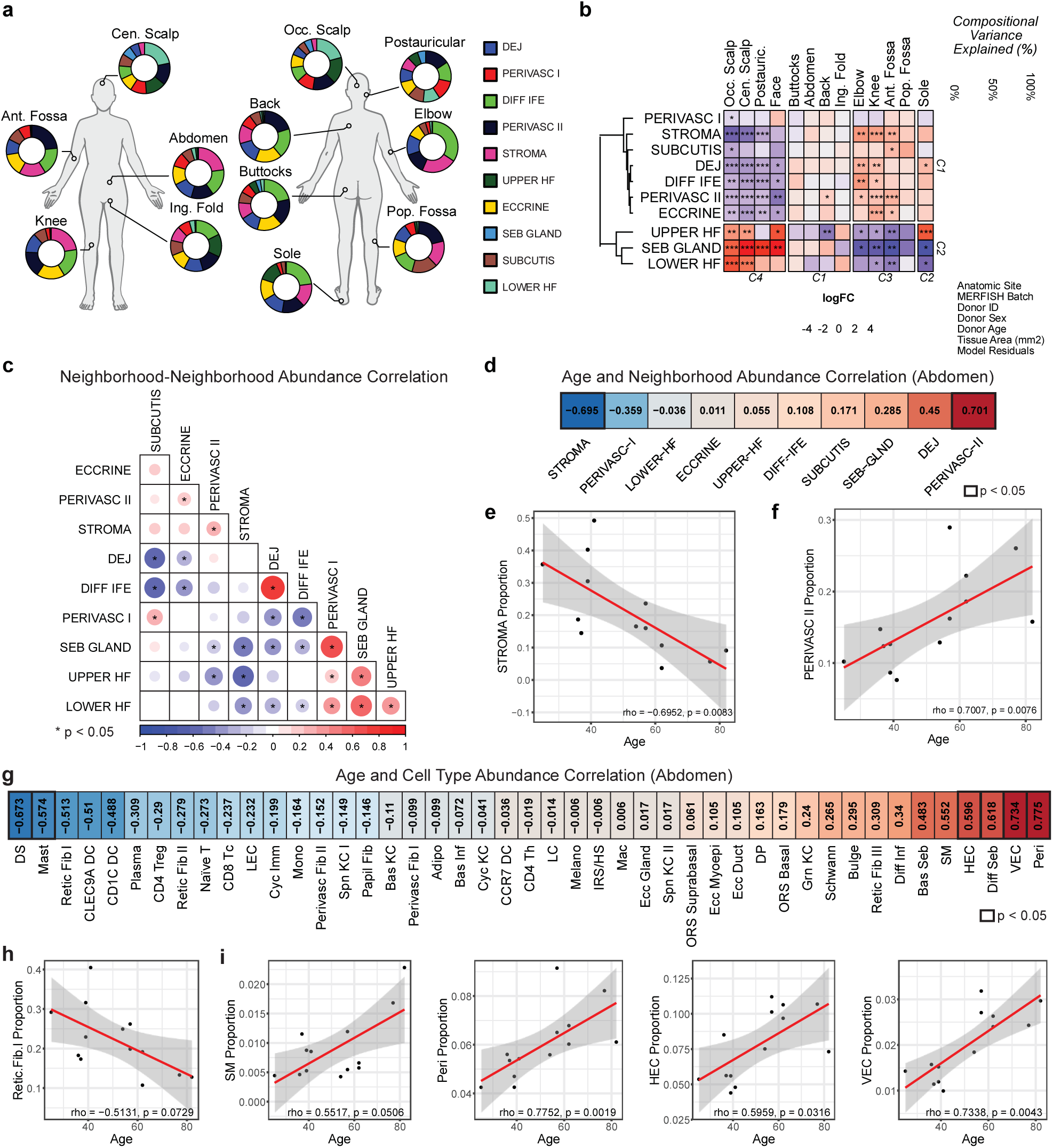
Neighborhood abundances vary across anatomic sites and during aging. **a.** Donut plots of neighborhood proportions at each site across N = 7 donors with 12 anatomic sites each. **b.** Top, bar plots of variance portioning showing contributions of each metadata to the variance in data. Bottom, clustered heatmap of neighborhood abundances across anatomic sites. -adjusted p < 0.1, *adjusted p < 0.05, **adjusted p < 0.01, ***adjusted p < 0.001, moderated t test. **c.** Clustered pairwise neighborhood abundance Spearman correlations across 13 abdomen samples (N = 13 donors). **d.** Ranked Spearman correlation coefficients (r) of neighborhood abundance with age from abdomen samples (n = 13 samples from N = 13 donors). **e.** Scatter plot of STROMA neighborhood proportion and age from N = 13 donors. Red line = linear model fit, gray area = 95% confidence interval. **f.** Same as (e) but for PERIVASC II proportion and age. **g.** Ranked Spearman correlation coefficients (r) of cell type abundance with age from abdomen samples (n = 13 samples from N = 13 donors). **h.** Scatter plot of Retic Fib I proportion and age from N = 13 donors. Red line = linear model fit, gray area = 95% confidence interval. **i.** Same as (h) but for additional cell type proportion and age: smooth muscle (SM), pericyte (Peri), high endothelial venule-like endothelial cells (HEC), and vascular endothelial cell (VEC). Red line = linear model fit, gray area = 95% confidence interval.

Though we observed distinct neighborhoods across human skin, all neighborhoods are contiguous and can interact across neighborhood boundaries. To assess patterns of cross-neighborhood coherence and potential neighborhood-neighborhood interactions, we performed pairwise correlation analysis of neighborhood abundances across all samples. These data demonstrated a tight linkage of the DEJ and DIFF IFE (ρ = 0.74) consistent with these two neighborhoods working together to drive epidermal thickness as well as a correlation between STROMA and PERIVASC II (ρ = 0.26) (**Figure 4c**). Meanwhile, SEB GLAND exhibited high correlation with both UPPER HF and LOWER HF as well as PERIVASC I. In contrast, ECCRINE exhibited low correlation with every other neighborhood except for PERIVASC II, consistent with more independent or skin-extrinsic regulation (**Figure 4c**). These analyses suggest that some neighborhoods may be more biologically interactive with one another than others.

Finally, our dataset also offered a unique opportunity to assess how aging affects neighborhood and cell type abundance dynamics. Since anatomic site is a major determinant of neighborhood abundance, we restricted this analysis to abdomen, from which we obtained samples from 13 donors. Correlating neighborhood abundance across age highlighted significant correlations of STROMA (ρ = -0.70) and PERIVASC II (ρ = 0.70), with no other neighborhoods exhibiting significant correlation (**Figure 4d**). Given the correlation of these two neighborhood abundances (**Figure 4c**), aging appears to be associated with a shift from STROMA to PERIVASC II abundance. Cell types associated with this shift include a reduction of Retic Fib I (prominent constituent of STROMA) and increases in HEC, VEC, Peri, and SM (prominent constituents of PERIVASC II) (**Figure 4e**), highlighting cellular changes during aging that potentially affect neighborhood function. While prior studies have shown reduction in gene expression by fibroblast subsets with age in the inguinal fold, here we observe that at the abdomen, numbers of Retic Fib I are reduced. In sum, we identified 10 multicellular neighborhoods that define the cellular spatial organization in skin, including an immune-enriched PERIVASC I neighborhood, neighborhood-neighborhood relationships, and neighborhoods that are dynamic across anatomic sites and during aging.

### Neighborhoods define distinct spatial cell-cell communication networks

We next mapped spatially resolved ligand–receptor **(L–R)** interactions within each MERFISH neighborhood, revealing distinct intercellular signaling patterns using CellChat analysis of MERFISH expression data **(Table S5)**. Given high diversity of the PERIVASC I neighborhood and its enrichment of immune populations, we examined this neighborhood further (**Figure 3c-d** and **Table S5**). Within PERIVASC I, top interacting cell types included VECs, CD4+ Tc, CD8+ Tc, and Perivasc Fib I/II (**Figure 5a**). To extend beyond MERFISH panel genes and mitigate data sparsity, we simulated neighborhoods using our integrated scRNA-seq reference and repeated L–R inference with CellChat (**Extended Data Figure 8a** and **Methods**). Using this approach, we identified top pathways enriched within PERIVASC I, which included MIF, midkine (MK), CXCL/CCL chemokines, and TNF (**Figure 5b**). Expanding these analyses across all neighborhoods in scRNA-seq data revealed that PERIVASC I and ECCRINE had the most inferred interactions (**Extended Data Figure 8a-b**). We also identified the top unique and shared L-R pathways across neighborhoods (**Extended Data Figure 8c**). Consistent with the heterotypic interactions among immune and stromal cells, PERIVASC I exhibited the highest number of unique pathways, while most other neighborhoods shared common signaling pathways (**Extended Data Figure 8c**). Top L-R pairs in the PERIVASC I neighborhood included CXCL12-CXCR4, MDK-CD74, MIF-CD74, and PTN-NCL as highest predicted stromal-to-immune interactions (**Figure 5c**). Conversely, immune-to-stromal interactions involved PPIA-BSG, NAMPT-ITGA5+ITGB1, and TNF-TNFRSF1A (**Figure 5d**). Scoring MERFISH panel L–R pairs, such as CXCL12–CXCR4, between anchoring cell types and their nearest neighbors confirmed perivascular co-expression, indicating that immune–stromal crosstalk sustains the PERIVASC I neighborhood (**Figure 5e** and **Methods**).

**Figure 5.**
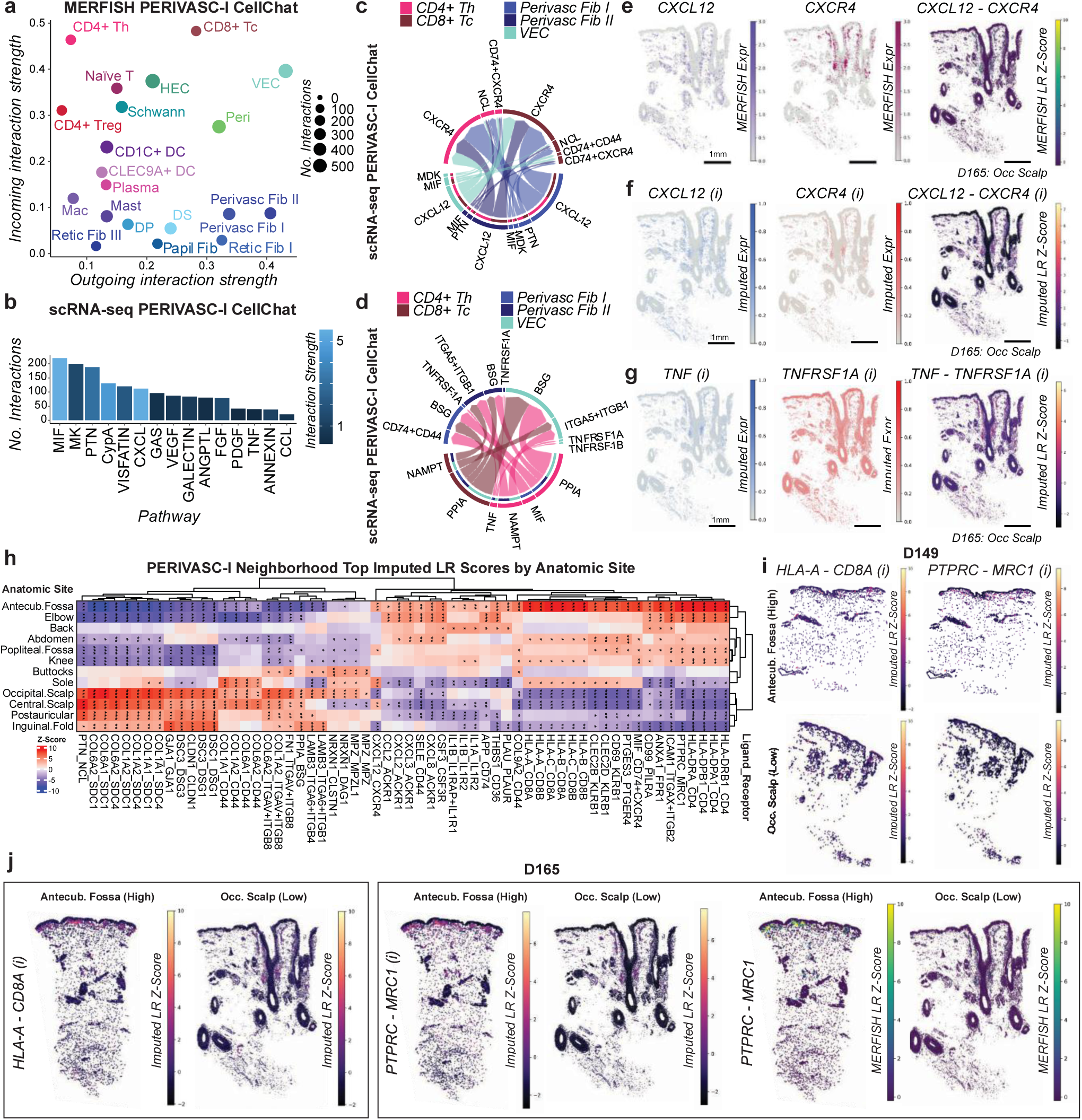
Neighborhoods define cell-cell communication within and across sites. **a.** Scatter plot of incoming and outgoing interaction strengths per cell type in PERIVASC-I neighborhood predicted by CellChat on MERFISH data. **b.** Highest predicted active pathways in PERIVASC-I neighborhood from scRNA-seq data. **c.** Circos plot of top stromal to immune ligand-receptor communication pairs in scRNA-seq of cell types in PERIVASC-I neighborhood. **d.** Circos plot of top immune to stromal ligand-receptor communication pairs in scRNA-seq of cell types in PERIVASC-I neighborhood. **e.** *CXCL12* and *CXCR4* expression in MERFISH data, and ligand-receptor (LR) score from D165 occipital scalp (see Methods). **f.** Imputed (i) *CXCL12* and *CXCR4* expression, and imputed ligand-receptor (LR) score from D165 occipital scalp (see Methods). **g.** Imputed (i) *TNF* and *TNFRSF1A* expression, and imputed ligand-receptor (LR) score from D165 occipital scalp. **h.** Heatmap of top ligand-receptor (LR) scores at each site in the PERIVASC-I neighborhood. *adjusted p < 0.05, **p < 0.01, ***p < 0.001, moderated t test. **i.** Imputed (i) ligand-receptor (LR) scores from D149 of indicated LR pairs at high (antecubital fossa) and low (occipital scalp) scoring sites and **j.** from D165. Non-imputed MERFISH LR scores are also shown for *PTPRC MRC1* (a pair included in MERFISH panel).

We next visualized the spatial expression of the L-R pairs predicted by scRNA-seq that were not in our MERFISH panel using Tangram imputation (**Extended Data Figure 9a** and **Methods**).^37^ We first verified that imputed values correlated with ground truth MERFISH expression patterns, focusing on accurate expression of well-known marker genes for KCs, fibroblasts, eccrine glands, and endothelial cells, in addition to L-R pairs included in our MERFISH panel (**Figure 5e-f** and **Extended Data Figure 9b**). This approach enabled visualization of additional L–R pairs absent from the MERFISH panel, such as TNF–TNFRSF1A, which showed high PERIVASC I, DEJ, and SUBCUTIS scores consistent with scRNA-seq results (**Figure 5g** and **Extended Data Figures 8c and 9c**). Imputation also allowed us to examine molecular variation across anatomic sites. Differential L–R scoring identified the top interactions within PERIVASC I at each site (**Figure 5h, Table S5,** and **Methods**). In the antecubital fossa, we observed elevated expression of both MHC class I (HLA-A/B/C/E) and II (HLA-DRA/DMA/DQA1) co-expressed with CD8 and CD4, potentially enhancing T cell retention, alongside chemokine upregulation (CXCL2/3/8/12, CCL2) and MIF–CD74+CD44 co-expression, which may further drive T cell recruitment and activity (**Figure 5h-i** and **Table S6**). In the DEJ, higher expression of ECM components (COL1A1/2, COL6A1/2, COMP) with their cognate receptors was consistently detected in sole, knee, and elbow across donors in both MERFISH and imputed data compared with other sites, suggesting a molecular basis for increased epidermal thickness (**Extended Data Figure 9d-h** and **Table S6**). Together, MERFISH directly resolved top cell–cell interactions within our panel, while imputation expanded analyses transcriptome-wide, revealing site-specific molecular programs that may underlie anatomical variation in cell composition, including enhanced immune presence at flexural sites.

### Skin neighborhood remodeling in disease

To determine how microanatomic neighborhood dynamics are altered in disease, we curated publicly available spatial datasets (10X Visium) from normal human skin and diseased tissues, including the inflammatory skin diseases atopic dermatitis (AD), psoriasis (PP), and hidradenitis suppurativa (HS), and skin cancers basal cell carcinoma (BCC) and squamous cell carcinoma (SCC) (**Figure 6a**). After filtering poor capture spots, we integrated 142,515 capture spots from 81 tissue samples (44 normal skin, 7 AD, 4 PP, 10 HS, 8 BCC, and 8 SCC) from 63 donors (**Figure 6b** and **Methods**).

**Figure 6.**
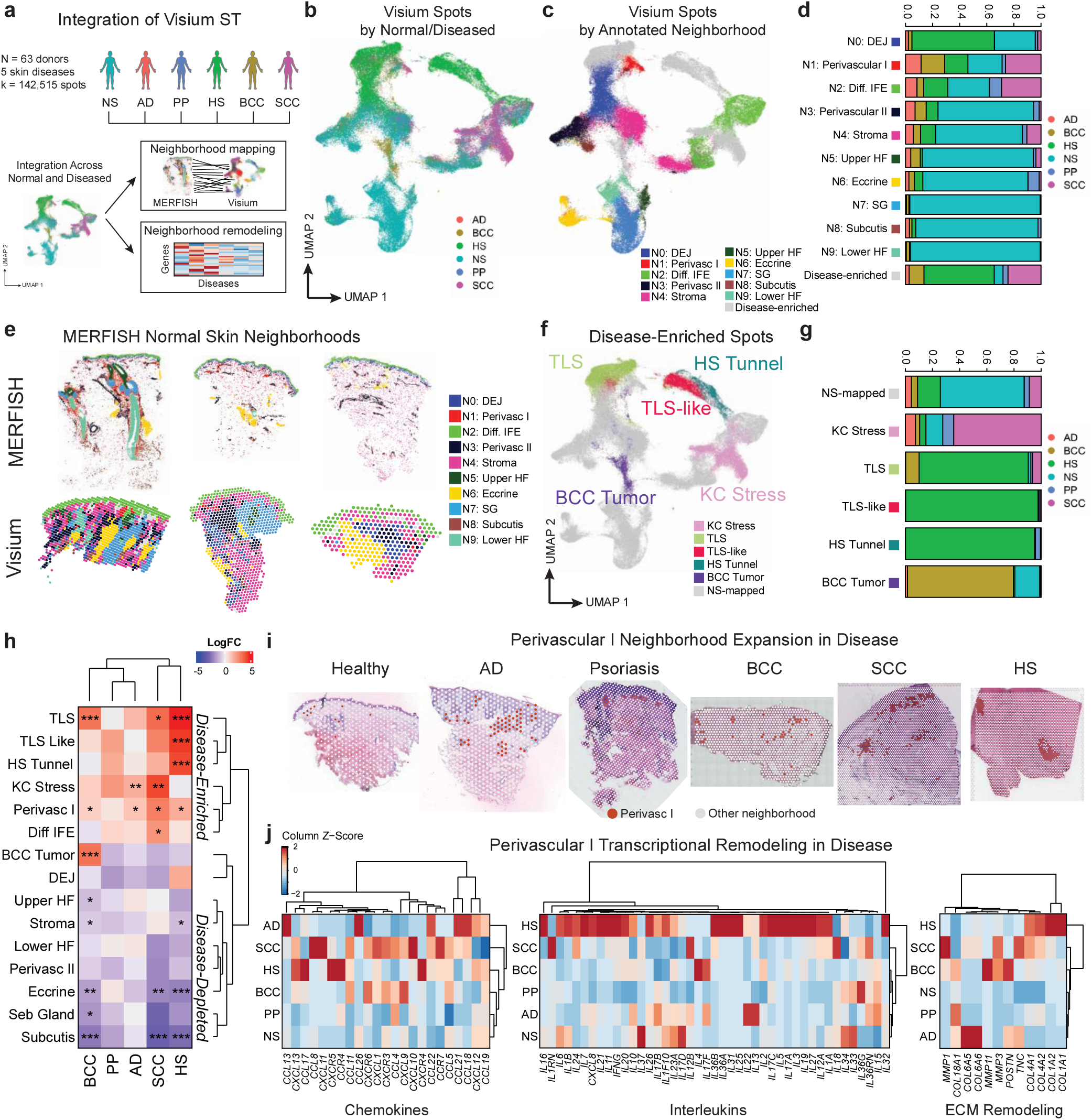
Neighborhoods define microanatomic remodeling in human skin disease. **a.** Workflow for Visium data integration and downstream analysis encompassing neighborhood mapping and neighborhood transcriptional remodeling. NS = normal skin, AD = atopic dermatitis, PP = psoriasis, HS = hidradenitis suppurative, BCC = basal cell carcinoma, SCC = squamous cell carcinoma. **b.** UMAP of integrated Visium spots across all samples labeled by normal and diseased tissue state (n = 142,515 spots). **c.** Same UMAP as (B) labeled by annotated the MERFISH neighborhoods they resemble the most. **d.** Bar plots of tissue state proportions comprising each mapped MERFISH neighborhood. **e.** Spatial plots of neighborhoods in representative samples of each tissue state. **f.** Same UMAP as (B) and (C), labeled by annotated for disease-enriched neighborhoods. **g.** Bar plots of tissue state proportions comprising each disease-enriched neighborhood. **h.** Heatmap of neighborhood abundances of diseased tissues relative to normal skin. *adjusted p < 0.05, **adjusted p < 0.01, ***adjusted p < 0. 001, moderated t test. **i.** Perivascular I neighborhood cluster highlight across representative samples of each tissue state. **j.** Heatmap of chemokine (left), interleukin (middle) and ECM remodeling (right) gene expression in Perivascular I spots across tissue states.

Given that the 55µm capture spot size in 10X Visium contains tens of cells, we reasoned that each spot encompasses at least a portion of a multicellular neighborhood, and that unbiased clustering of spots would recover groups of spots comparable to neighborhoods (or components thereof) in MERFISH data. Unbiased clustering identified 20 clusters that largely exhibited features of epithelial, immune, and stromal compartments both transcriptionally (via top differentially expressed genes) and spatially (via visualization of underlying histology) (**Extended Data Figure 10a-b**). We next sought to map Visium clusters to the 10 MERFISH neighborhoods we had previously identified. First, we focused on clusters enriched in normal skin: C0 expressed mostly fibroblast-related genes and resembled the DEJ and STROMA neighborhoods; C12 and C18 highly expressed eccrine gland markers resembling the ECCRINE neighborhood; C5 and C11 expressed sebaceous markers, resembling the SEB GLAND neighborhood; C9 expressed hair-follicle associated keratins, resembling the LOWER HF neighborhood; C16 expressed epithelial differentiation markers, resembling the UPPER HF neighborhood; C2 and C6 expressed ECM and fibroblast genes resembling the STROMA neighborhood; C15 contained a mix of vasculature, immune, and fibroblast markers highly resembling the PERIVASC I neighborhood; C7 expressed smooth muscle and lymphatic EC markers, resembling the PERIVASC II neighborhood; C17 expressed adipocyte markers, resembling the SUBCUTIS neighborhood; and C4 and C10 expressed epidermal differentiation markers highly resembling the DIFF IFE neighborhood (**Extended Data Figure 10a-b** and **Table S7**). Scoring each capture spot using neighborhood signatures derived from top expressed genes within each neighborhood further confirmed these neighborhood annotations of Visium spots (**Figure 6c-e, Extended Data Figure 10c,** and **Methods**). To additionally assess accuracy of spot cluster annotations, we deconvolved cell type composition of spots with cell2location^38^ using our compiled scRNA-seq dataset as a reference. Cell2location results were consistent with inferred cell type composition using differential gene expression, demonstrating multiple distinct cell types within each cluster that mirrored MERFISH neighborhood composition (**Extended Data Figure 10d**). Thus, Visium data captured key aspects of our MERFISH neighborhoods across normal skin tissues and provided a basis for examining disease-associated clusters.

We next examined disease-associated clusters that were enriched in disease and further annotated them based on differentially expressed genes within the cluster (**Extended Data Figure 10a and 11a**). We identified four disease-enriched clusters (C1, C3, C4, and C15) that exhibited increased abundance in more than one disease (**Extended Data Figure 11a**). The KC Stress neighborhood (C1) was most expanded in SCC and encompassed SCC tumor areas marked by stress-related keratins *KRT6A/B/C* and *S100A8/9* (**Extended Data Figure 11b** and **Table S7)**. Notably, they were also present in the epidermal layers of AD and BCC (in non-tumor overlying epidermis), indicating high keratinocyte reactive stress in these diseases. Notably, as described above, C4 and C15 both mapped strongly to normal skin neighborhoods. C4, which resembled DIFF IFE in normal skin, was expanded in SCC, HS, and AD and was marked by keratinization genes, which is consistent with reactive differentiation in response to disease-associated stimuli such as inflammation (**Table S7**). C15, in addition to its resemblance to the normal skin PERIVASC I neighborhood, was expanded in SCC, BCC, HS, and AD, was marked by increased expression of T cell (*TRAC*/*TRBC1*/*TRBC2*/*CD2*) and perivascular fibroblast (*CCL19*) markers, and chemokines (*CCL5*/*CCL13*/*CCL18*/*CXCL9*/*CXCL10*) (**Extended Data Figure 11c**). C3 was enriched in SCC, BCC, HS, and AD and was marked by high plasma cell marker expression (*IGLC1*/*IGHM*/*JCHAIN*/*CD79A*) and chemokines (*CXCL12*/*13*), and it corresponded to histological tertiary lymphoid structure (TLS) structures (**Extended Data Figure 11d**). Among clusters uniquely expanded in a single disease, C13 and C19 were specific to BCC and corresponded to BCC tumor areas, marked by *PTCH1/2*, *SMO*, and *EPCAM* in C13 (**Extended Data Figure 11a** and **11e**). C19 was a minority cluster with minimal gene expression differences (**Table S7**). C8, C14, and C0 additionally favored HS (**Extended Data Figure 11a**). C14 corresponded with HS-associated epithelial tunnels, a hallmark of advanced disease (**Extended Data Figure 11b** and **11f**). C8 was similar to C3 in expressing TLS-like genes but at lower levels, and C8 exhibited spatial proximity to the epithelial tunnels, suggesting a potential response to HS-specific tunnels (**Extended Data Figure 11f-g**). Finally, C0 was notable for its presence in normal skin and high DEJ and STROMA neighborhood scores, and its expansion in HS may indicate potential mechanisms by which some HS-specific features may arise, such as epithelial tunnels that receive increased stromal signaling. Together, these data enabled detailed annotation of Visium spots enriched in disease (**Figure 6f-g**), and highlight both the shared (e.g., keratinocyte stress, immune infiltration) and unique (e.g., BCC tumor cells, HS epithelial tunnels) transcriptional and spatial features across skin diseases, supporting the concept of spatially compartmentalized pathologic responses.

We next asked how these Visium-annotated neighborhoods shift during disease. Differential abundance analyses highlighted the Perivascular I neighborhood spots as the most broadly dynamic disease-enriched neighborhood, with expansion in four of five diseases sampled (**Figure 6h-i**). We then examined gene expression of biologically relevant processes within this neighborhood across normal and diseased tissues. The Perivascular I neighborhood exhibited increases in chemokines, cytokines (interleukins), and ECM factors in diseased skin compared to normal skin, suggesting that it is both a key spatial domain for immune recruitment and activity, and a potential target for therapeutic modulation (**Figure 6j**). Collectively, our pan-skin disease integrated analysis of Visium data suggests that skin disease is associated with both expansion and transcriptional remodeling of normal skin neighborhoods, with identification of the Perivascular I neighborhood as an immunomodulatory spatial domain. Since inflammatory skin disease results in excessive tissue inflammation and cancer results in an inadequate immune response, focused efforts to modulate perivascular cellular dynamics could yield improved therapeutic targeting in inflammatory skin disease and skin cancer.

## DISCUSSION

The spatial organization of human skin underlies its form and function both within the tissue microenvironment and across the body plan to achieve its many critical roles, including barrier establishment and immune surveillance. Here, we generated an adult human skin MERFISH atlas to localize 45 cell types across 15 anatomic sites with a unique donor-matched sampling strategy, defining homeostatic spatial neighborhoods and how disease can arise from their disruption. This comprehensive resource reveals insights into skin cellular organization at multiple levels of granularity and can be mined for further exploration beyond the questions considered here.

### MERFISH skin optimization enables single-cell deconvolution comparable to scRNA-seq

We performed extensive optimization of human skin tissue preparation to generate this MERFISH atlas, setting new standards in terms of resolution and scale, coupled with careful custom gene panel design that enabled successful deconvolution of 45 cell types. As spatial technologies improve to address current tradeoffs in gene coverage and sensitivity (breadth vs. depth), our data will serve as a benchmark for future studies in terms of transcript sensitivity and cell type granularity. Attesting to broader impacts on the spatial biology field, our data could further accelerate improvements in spatial analysis tools, including cell segmentation algorithms that represent a major challenge in imaging-based spatial methods.

Our atlas enabled a spatial census of cell types previously shown in scRNA-seq studies, which we harmonized into a reference integrated scRNA-seq atlas for cross-referencing cell type annotations. In addition to achieving cross-modal spatial-scRNA coherence, MERFISH additionally captured cells such as sebocytes and adipocytes that are notoriously difficult to recover during tissue dissociation-based assays. Improved cell type coverage allowed for highly accurate analyses of cell type abundances free from dissociation biases that limit similar efforts in scRNA-seq studies, while still accounting for donor-level variation. Cell type abundance patterns recapitulated body axes with remarkable consistency, including centrifugal (trunk to extremities), anterior-posterior (extensor to flexural), and craniocaudal (scalp to buttocks/sole) coordinates. In addition to expected patterns that were previously identified in scRNA-seq (e.g., increased Spn KC II and Retic Fib II in sole), and higher abundance of hair follicle-associated cell types in scalp, we identified immune cell enrichment in the flexural areas, and papillary fibroblast enrichment in the sole, knee, and elbow, which correlated with overall epidermal thickness.

We localized fibroblast scRNA-seq subpopulations that were difficult to probe with traditional immunofluorescence or smFISH due to overlapping markers, in addition to the extensive repertoire of immune cells that populate the skin, which congregated almost exclusively around the vasculature. MERFISH fibroblast subpopulations included well-known Papillary and Reticular (Retic Fib I) populations, but additionally included sole-enriched Retic Fib II, whose previous spatial location was unknown, two perivascular (superficial Perivasc Fib I and deeper Perivasc Fib II), perineural fibroblasts that co-localize with nerve bundles (Schwann cells), and hair follicle-associated DS and DP-like populations. The distinct locations suggest specific subpopulation roles, deepening our understanding of the function of heterogeneous fibroblast subsets. For example, while papillary fibroblasts are well-established epidermal signalers, our analyses establish their clear correlation with epidermal thickness, including in the sole, where Retic Fib II cells are also located but scattered throughout the dermis, suggesting additional structural roles for Retic Fib II. Given recent in-human studies demonstrating the ability of sole fibroblasts to induce increased epidermal thickness,^13^ further clarification on whether Papillary or Retic Fib II (or both) can induce thicker skin could refine this promising cellular therapy. Since we captured relatively few anagen-phase hairs in our tissue sections, and even fewer hair bulbs where bona fide DP cells should reside, we were surprised to capture the DP-like population that expresses canonical DP markers such as *CRABP1* and the dermal sheath marker *LRRC15.*^39,40^ This population localized mostly around hair follicles, but further studies will be required to establish its role. We speculate it may contain previously described hair follicle dermal stem cells (hfDSCs) that serve as a reservoir of DP and DS cells.^41^

### Neighborhoods provide a cellular blueprint for reconstructing human skin spanning the molecular to anatomic scale

The 45 distinct cell types identified by MERFISH were spatially organized into 10 multicellular neighborhoods, each associated with unique cell-cell interactions—that represent core building blocks of a fully functional skin tissue. Neighborhoods varied across anatomic site and age, demonstrating a shift from the STROMA to PERIVASC II neighborhood during aging, corresponding with decreasing Retic Fib I and increasing VEC/HEC/Pericyte abundances. Given the known loss of collagen production with age,^42^ future studies could focus on modulating or replacing Retic Fib I cells (possibly through cellular therapy) for improving skin integrity during aging. In addition to expected HF-associated neighborhood enrichment at hair-dense sites such as the scalp, the sole was surprisingly enriched in the UPPER HF neighborhood, highlighting infundibular-like features of the plantar epidermis. Within neighborhoods, we identified both direct spatial ligand-receptor interactions from our MERFISH data and imputed transcriptome-wide interactions using an integrated scRNA-seq reference. Broadly, these neighborhoods suggest a requirement of homeostatic cell-cell interactions for maintaining specific cell types in the skin. As *ex vivo* tissue engineering efforts, including human skin organoids, strive to incorporate immune components, our data provide a blueprint for reconstructing such neighborhoods, including PERIVASC I cellular components and molecular factors (e.g., L-R pairs upregulated in the antecubital fossa). In addition, adnexal structures that have yet to be fully recapitulated *ex vivo*, such as terminal hair follicles and sweat glands, could be built through UPPER/LOWER HF and SEB GLAND neighborhood and ECCRINE neighborhood reconstruction, respectively. These efforts to make fully functional skin *ex vivo* could unlock the next generation of therapeutic testing platforms that aim to combat disease and aging and augment regeneration.

### A perivascular neighborhood hosts homeostatic immune surveillance in the skin

Given the congregation of most immune cells, including DCs and T cells, in the PERIVASC I neighborhood, this neighborhood may well be considered the skin-associated lymphoid tissue (SALT), analogous to mucosa-associated lymphoid tissue (MALT) in tissues such as the oral cavity (tonsils) or small intestine (Peyer’s patches). Like other lymphoid tissues, this neighborhood also contains a key stromal cell, *CCL19*+ Perivasc Fib I, that resemble the fibroblast reticular cells of secondary lymph nodes, and we speculate that it serves a similar immune-organizing role.^43^ Furthermore, several predicted interactions between immune cells and Perivasc Fib I suggest that homeostatic immune cytokine signaling such as TNF-a may be required to maintain this stromal population, and reciprocally, fibroblasts may help maintain immune recruitment and residence (e.g., via CCL19-CCR7 and CXCL12-CXCR4). Additional anatomic site variation of immune and stromal cell abundances and co-expression of crosstalk molecules, such as the increase in MHC Class I/II with CD4/CD8 receptors in the antecubital fossa, hint at adaptive inflammatory setpoints at distinct body sites. While these may have evolutionarily arisen to protect more vulnerable locations, they potentially lower the threshold for chronic inflammation and disease susceptibility. Follow-up work could focus on such hypotheses.

### Spatially compartmentalized neighborhoods are dysregulated in skin disease

Neighborhood dynamics are disrupted during skin disease, as indicated by our analyses of previously published Visium data encompassing healthy skin and five skin diseases. Recent studies of inflammatory skin diseases have highlighted the presence of *CCL19*+ “pro-inflammatory” fibroblasts, observed in atopic dermatitis, psoriasis, hidradenitis suppurativa, and prurigo nodularis.^44–46^ Here, we map this population predominantly to the PERIVASC I neighborhood, which undergoes pathogenic immune expansion and remodeling during disease. The concept of induced SALT (iSALT) involving perivascular aggregates of DCs and T cells in mouse models of contact hypersensitivity has previously been shown,^47,48^ which highly resembles the PERIVASC I neighborhood. Given the presence of *CCL19*+ fibroblasts in both homeostasis and disease, these and other stromal cells may additionally be an important component of iSALT. Indeed, we and others have shown that targeting pro-inflammatory fibroblasts alone^49–51^ is sufficient to decrease immune infiltration in mouse models of skin inflammation, highlighting their potential as therapeutic targets. Furthermore, fibroblasts are key components of tertiary lymphoid structures (TLSs) observed in chronic inflammation and cancer,^52^ which we observed in Visium data as a distinct disease-associated neighborhood. Whether the formation of TLSs represents one extreme end of an iSALT, and whether they require multiple fibroblast subtypes identified here, could be examined in future studies. Our Visium analyses further support the concept that immune and stromal interplay in PERIVASC I underlies pathogenic inflammatory responses, in addition to identifying other key neighborhoods that are remodeled in disease. Collectively, understanding how skin neighborhoods are constructed, preserved, and vary across the human body may be key to maintaining healthy skin throughout our lifespan.

### Limitations of the study

There are several technical limitations in our data. First, MERFISH and other imaging-based spatial transcriptomics suffer from sensitivity limits with lower sensitivity than scRNA-seq. Further, both sensitivity and gene panel size restrictions limit cell type granularity achievable with scRNA-seq. For example, we could not identify rare cell types such as ILCs and NK cells confidently due to lack of reliable detection of distinguishing markers. Neurons were also absent due to failure to detect RNA transcripts within individual axons. As an imaging-based platform, MERFISH relies on segmentation for assigning transcripts to cells, which can lead to segmentation “doublets,” where segmented cells are assigned transcripts from two or more neighboring cells, which we filtered from downstream analysis. We also observed areas of likely over-segmentation, whereby larger cells were segmented into multiple smaller cells. These cases further highlight opportunities for improved analysis of our dataset through the development and optimization of novel segmentation algorithms. While our study considered certain donor metadata factors such as age and gender, we could not account for how additional environmental factors, such as sun exposure or sun-protective practices, impact the analyses. Finally, MERFISH imputation from scRNA-seq data should be interpreted cautiously, as we could not generate ground truth matched validation of scRNA-seq data from MERFISH tissues. Despite these limitations, our data should serve as a foundation for future human skin biology and disease studies.

## RESOURCE AVAILABILITY

MERFISH data are available on Zenodo and on an interactive website to explore the data is available online at https://rstudio-connect.hpc.mssm.edu/humanskin-spatialcensus/. Code and data are provided in a Zenodo repository with this paper that will be made available upon publication.

## ACKNOWLEDGEMENTS

We thank members of the Ji laboratory for helpful discussions. This work was supported by the National Institutes of Health (NIH) K08CA263187 (A.L.J.) and T32AR082315 (A.H. and R.G.), and Damon Runyon Cancer Research Foundation Clinical Investigator Award (A.L.J.). This research was conducted with the support of the Biorepository and Pathology CoRE at Icahn School of Medicine at Mount Sinai, the Bioinformatics for Next Generation Sequencing (BiNGS) shared resource facility within the Tisch Cancer Institute at the Icahn School of Medicine at Mount Sinai, which is partially supported by NIH grant P30CA196521, and in part by a pilot grant from the NIAMS/NIH SBDRC at Mount Sinai P30AR079200 grant. This work was supported in part through the computational and data resources and staff expertise provided by Scientific Computing and Data at the Icahn School of Medicine at Mount Sinai and supported by the Clinical and Translational Science Awards (CTSA) grant UL1TR004419 from the National Center for Advancing Translational Sciences. Research reported in this publication was also supported by the Office of Research Infrastructure of the National Institutes of Health under award number S10OD026880 and S10OD030463. The content is solely the responsibility of the authors and does not necessarily represent the official views of the National Institutes of Health.

## AUTHOR CONTRIBUTIONS

P.R. and A.L.J. conceived the project. A.W., S.O., A.H., A.K., I.D.S., L.C., I.I., R.G., M.G.H., A.S., and S.M. performed experiments and analyzed data. P.S. and A.L.J. performed bioinformatic analysis. A.R., D.D., and D.H. built the web resource. B.U., M.K., P.T., J.L., S.G., and S.M. interpreted data. A.L.J. guided experiments and data analysis and interpretation. P.R. and A.L.J. wrote the manuscript with input from all authors.

## EXTENDED DATA FIGURES

**Extended Data Figure 1.**
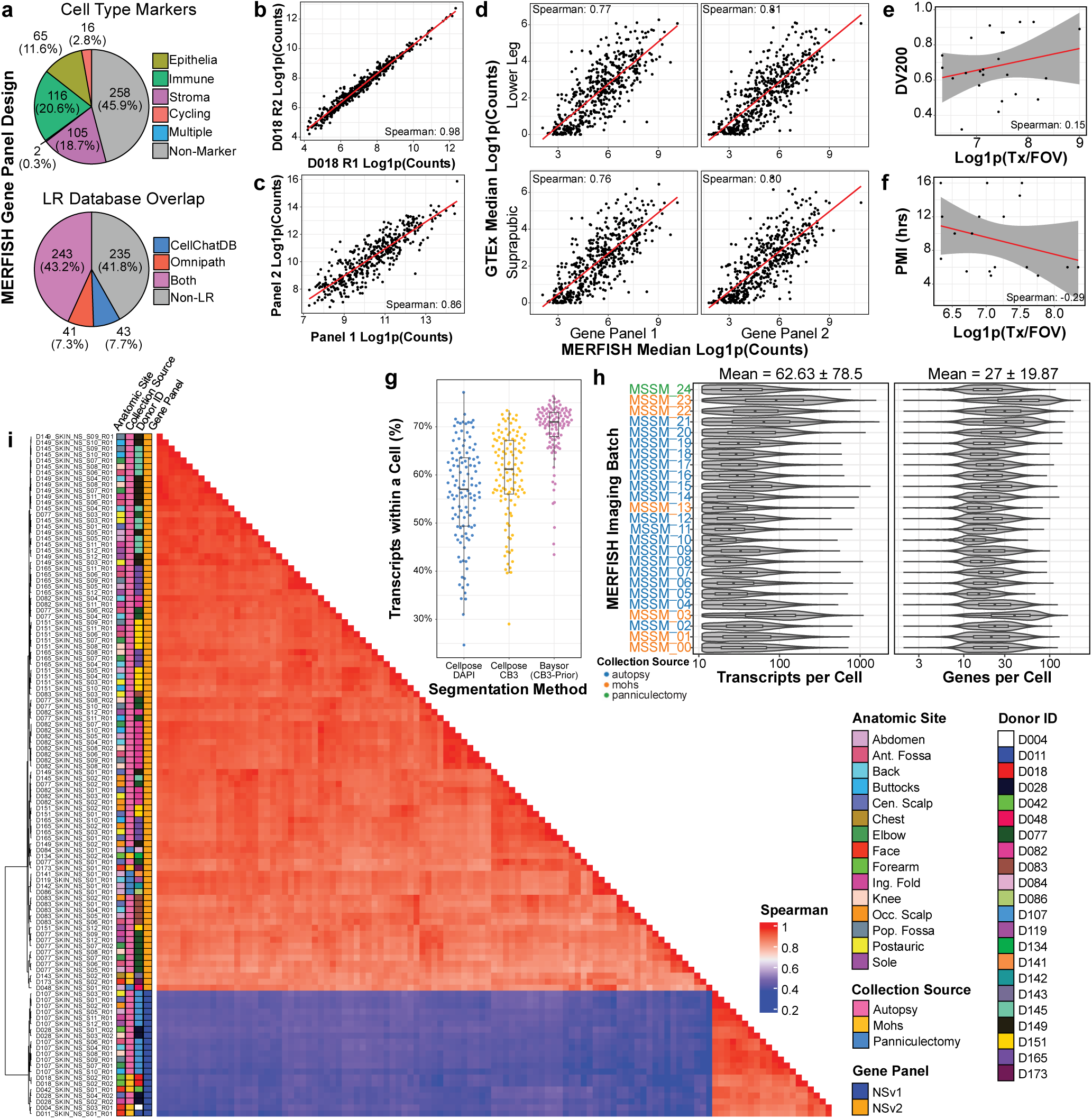
MERFISH achieves single-cell resolution profiling of adult human skin. **a.** Pie charts of cell type markers (top) and ligand-receptor (LR, bottom) gene distribution in MERFISH panels. **b.** Scatter plot of transcript counts across two replicate runs of adjacent sections of the same sample. Each point represents a gene (n = 500 genes). **c.** Scatter plot of summed transcript counts for all overlapping genes across all runs for each gene panel. Each point represents a gene (n = 434 genes). **d.** Scatter plots of median transcript counts in MERFISH runs for each panel and bulk RNA-seq data from GTEx human skin samples. **e.** Scatter plot of DV200 and transcript detection per field of view (Tx/FOV) for all MERFISH runs from FFPE samples (n = 23 runs). **f.** Scatter plot of post-mortem interval (PMI) and transcript detection per field of view (Tx/FOV) for all MERFISH runs featuring autopsy samples (n = 18 runs). **g.** Box plots of percent transcripts assigned to a cell with three segmentation approaches (see Methods) (n = 114 samples). Boxes display median and top and bottom quartiles of the data. **h.** Violin plots of transcripts per cell (post-filtering) for each MERFISH run (see Methods). **i.** Pairwise correlation of detected transcripts per sample, clustered (n = 114 samples).

**Extended Data Figure 2.**
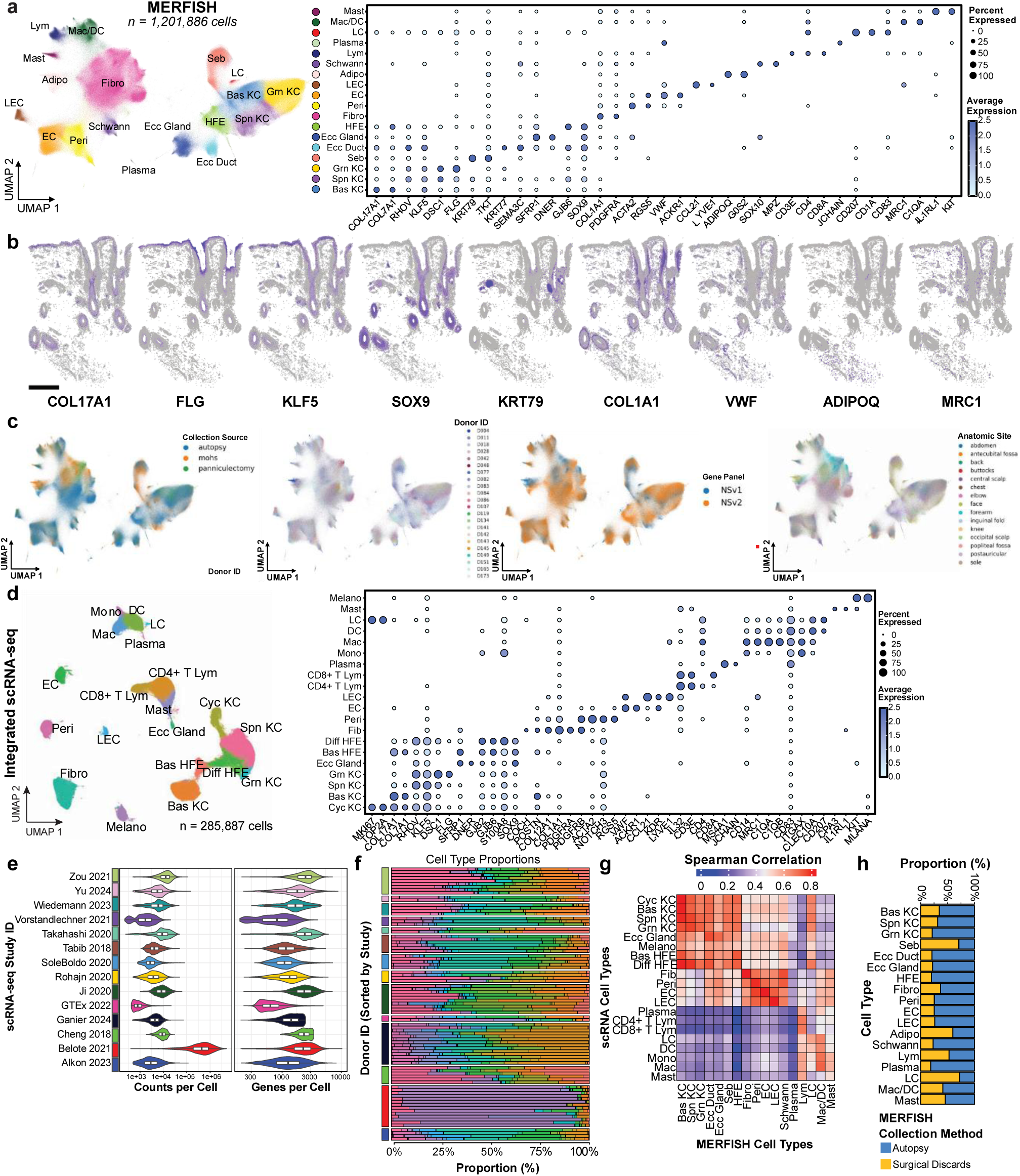
MERFISH and scRNA-seq deconvolve human skin at single-cell resolution. **a.** UMAP of integrated MERFISH data labeled broad cell types (left, n = 1,201,886 cells) and dot plot with expression of defining gene markers (right). **b.** Single-cell spatial localization of cell types labeled in (a) from 12 anatomic sites from a single donor (D165) profiled with MERFISH. Scale bar = 1mm. **c.** UMAPs of integrated MERFISH cells labeled by collection source (left), donor (middle), and gene panel (right). **d.** UMAP of integrated scRNA-seq data labeled by similar broad cell types as in (a) (left) and a dot plot with expression of defining gene markers (right). **e.** Violin plots of transcript counts per cell and genes per cell from 14 published scRNA-seq studies. **f.** Bar plots of proportion of each cell type per donor per study (N = 84 donors). **g.** Pairwise Spearman correlations of overlapping gene expression in MERFISH and scRNA-seq cell types. **h.** Bar plots of proportion of autopsy and surgical discard (fresh) samples per cell type in MERFISH data.

**Extended Data Figure 3.**
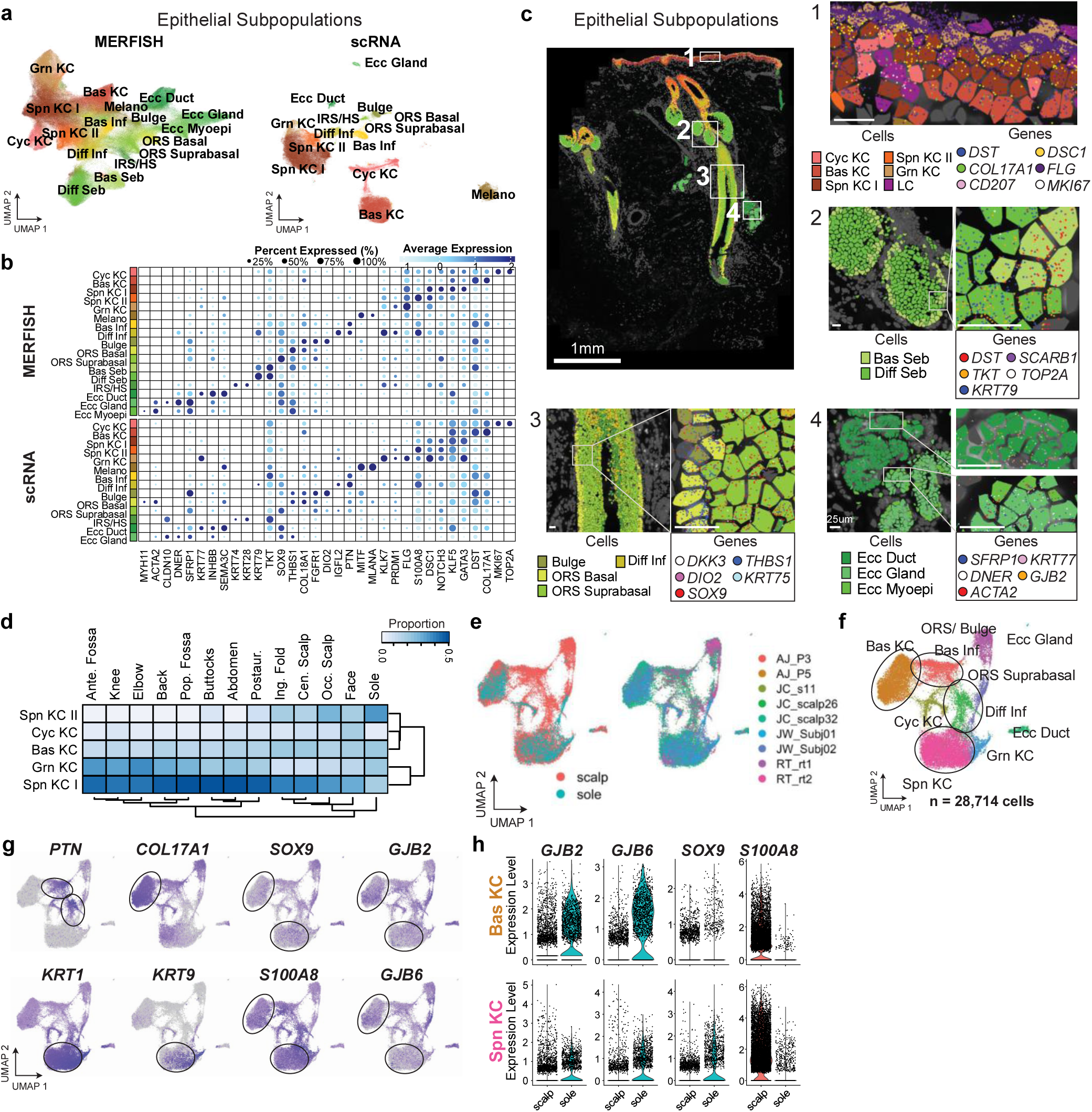
MERFISH identifies universal and site-enriched epithelial subpopulations. **a.** Integrated UMAPs (top) of MERFISH (n = 463,901 cells) and scRNA-seq (n = 139,691) epithelial subpopulations. **b.** Corresponding cell type marker dot plots of epithelial subpopulations in MERFISH (top) and scRNA-seq (bottom). **c.** Spatial localization of epithelial subpopulations in D165 postauricular sample with zoom-ins of subpopulations and top marker genes from the epidermis (1), sebaceous gland (2), hair follicle (3), and eccrine gland (4). Each dot represents a single transcript colored by its labeled gene. Scale bars = 25μm. **d.** Heatmap of proportion of IFE keratinocyte (KC) subpopulations per anatomic site in MERFISH data. Each row sums to 1. **e.** Integrated UMAPs of scalp and sole IFE keratinocytes isolated from scRNA-seq studies (N = 9 donors), labeled by site (left) and donor (right). **f.** Same UMAP as (e) but labeled by subpopulation. **g.** UMAP feature plots of expression of subpopulation markers. **h.** Violin plots of indicated gene expression in scalp and sole scRNA-seq Bas KC and Spn KC cells.

**Extended Data Figure 4.**
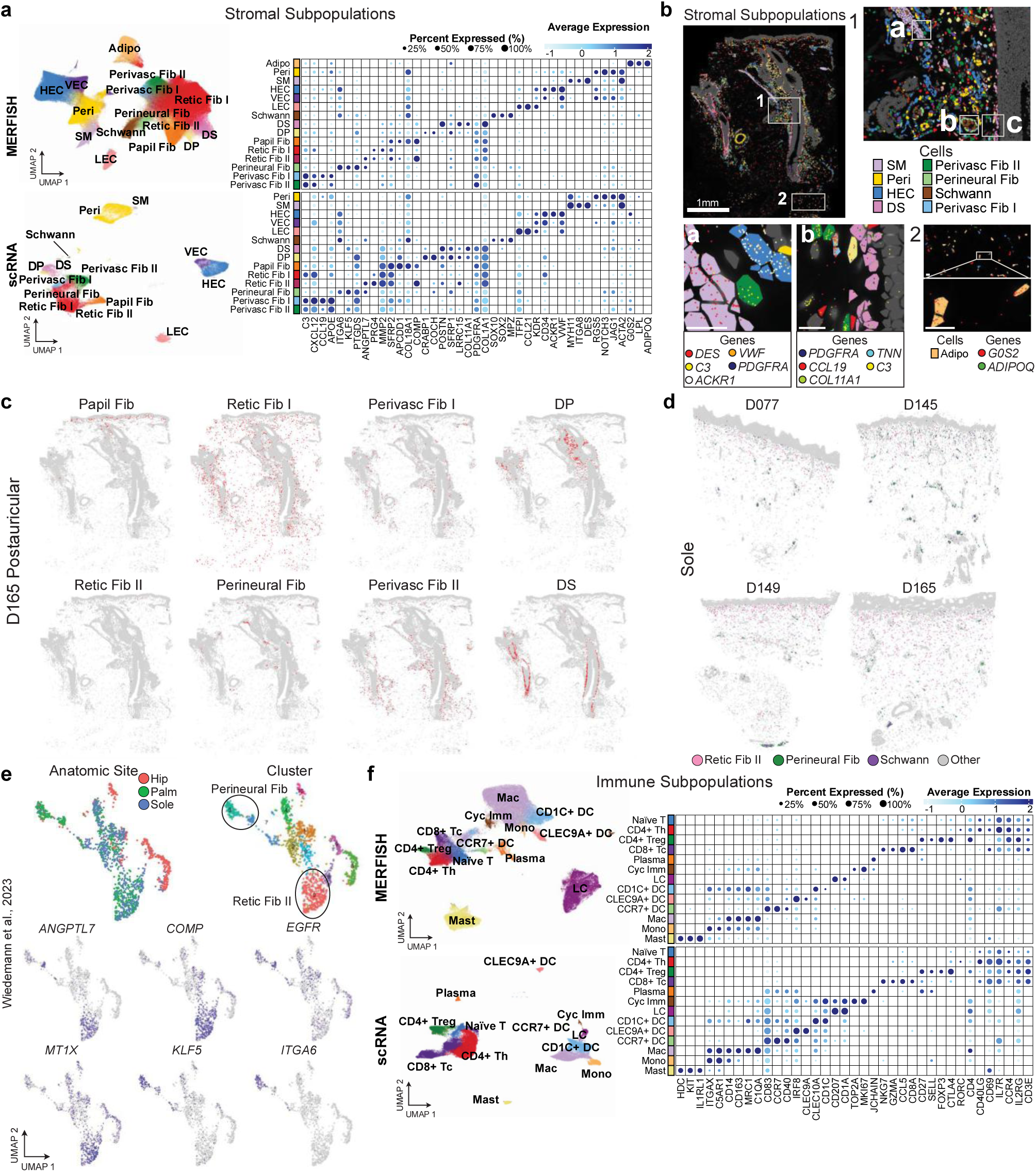
MERFISH catalogs stromal and Immune populations in adult human skin. **a.** Integrated UMAPs and corresponding cell type marker dot plots of MERFISH (top) (n = 526,010 cells) and scRNA-seq (bottom) (n = 60,119 cells) for stromal subpopulations. **b.** Spatial localization of stromal subpopulations in D165 postauricular sample with zoom-ins of subpopulations and top marker genes from the papillary dermis and superficial perivascular area (1); hair follicle (2) featuring SM of the arrector pili muscle (a), nerve bundle (b), and perifollicular area (c); and subcutis (3). Each dot represents a single transcript colored by its labeled gene. Scale bars = 25mm. **c.** Spatial highlights of fibroblast subpopulations in D165 postauricular sample. Red highlights each subpopulation, while gray indicates other cell types. **d.** Spatial highlights of Retic Fib II, Perineural Fib, and Schwann cells in sole samples from four donors. **e.** scRNA-seq analysis of hip, palm, and sole fibroblasts from Wiedemann et al. showing consistent expression of markers in Perineural Fib and Retic Fib II subpopulations. **f.** Integrated UMAPs and corresponding cell type marker dot plots of MERFISH (top) (n = 95,901 cells) and scRNA-seq (bottom) (n = 73,317 cells) immune subpopulations.

**Extended Data Figure 5.**
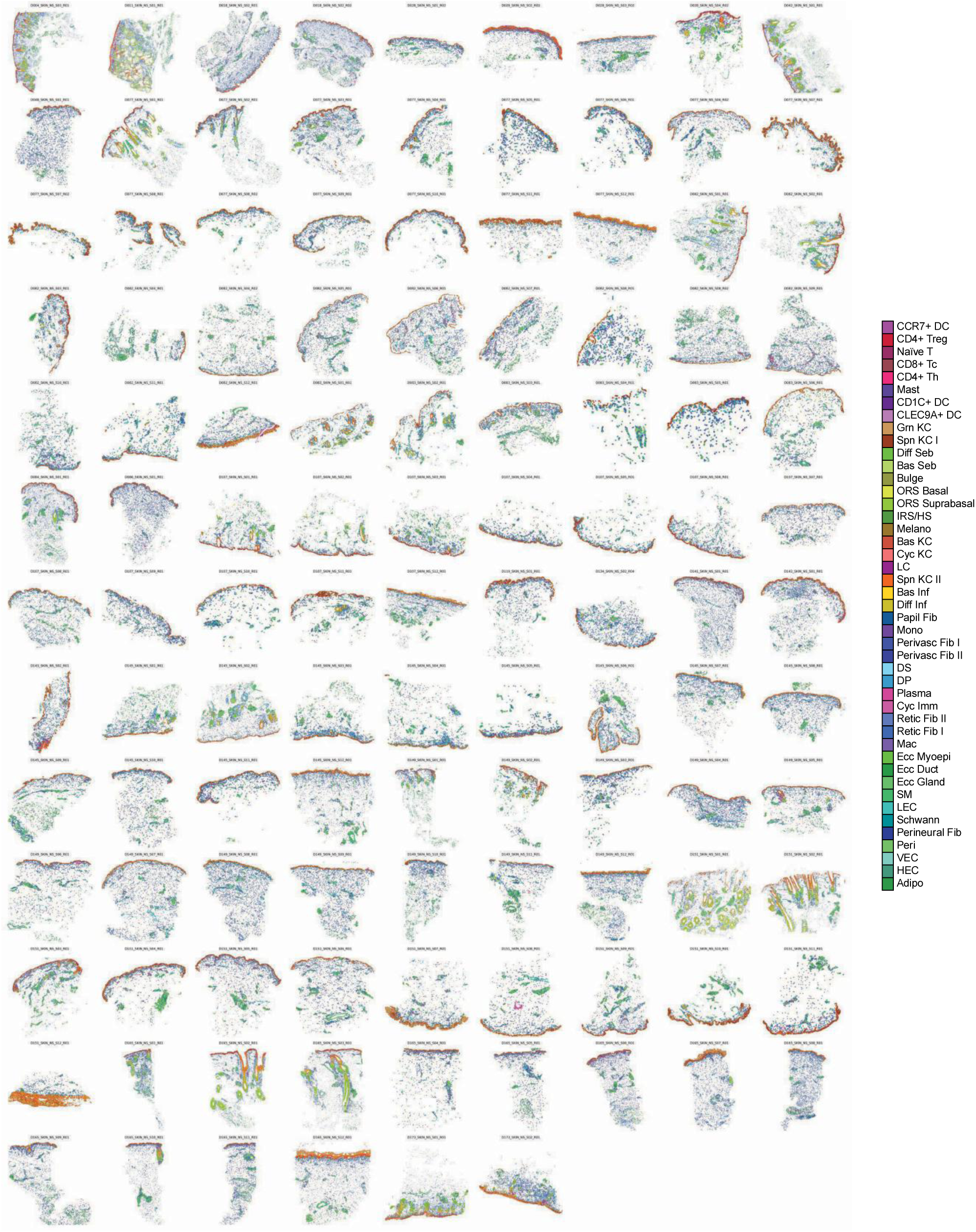
MERFISH identifies 45 subpopulations in human skin. Spatial localization of all subpopulations across all MERFISH samples (n = 114 samples).

**Extended Data Figure 6.**
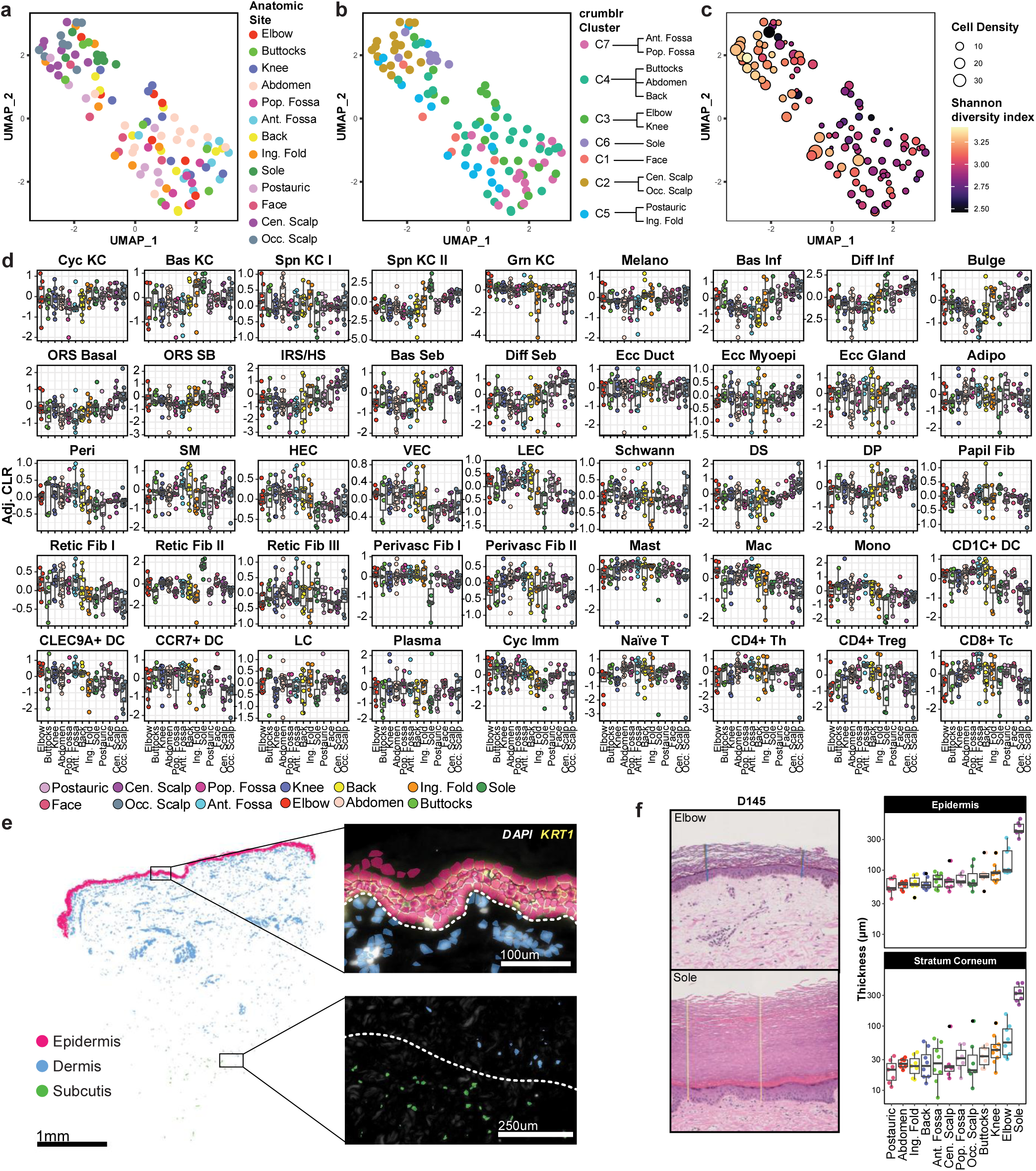
Cell type abundance variation underlies anatomic site differences. **a.** UMAP of samples using cell type abundances as features (n = 111 samples). **b.** Same as (a) but labeled by clusters determined by crumblr, as shown in Figure 2c. **c.** Same as (a) and (b) but labeled by cell density (point size) and colored by Shannon diversity index. **d.** Box plots of adjusted centered log ratios (CLRs) for cell types across sites (n = 111 samples). Boxes display median and top and bottom quartiles of the data. **e.** Schematic of manual annotation of epidermis, dermis, and subcutis compartments for all MERFISH samples. **f.** Left, example epidermal and stratum corneum thickness measurements from histology. Right, ranked box plots of average thickness of epidermis and stratum corneum across sites. Boxes display median and top and bottom quartiles of the data.

**Extended Data Figure 7.**
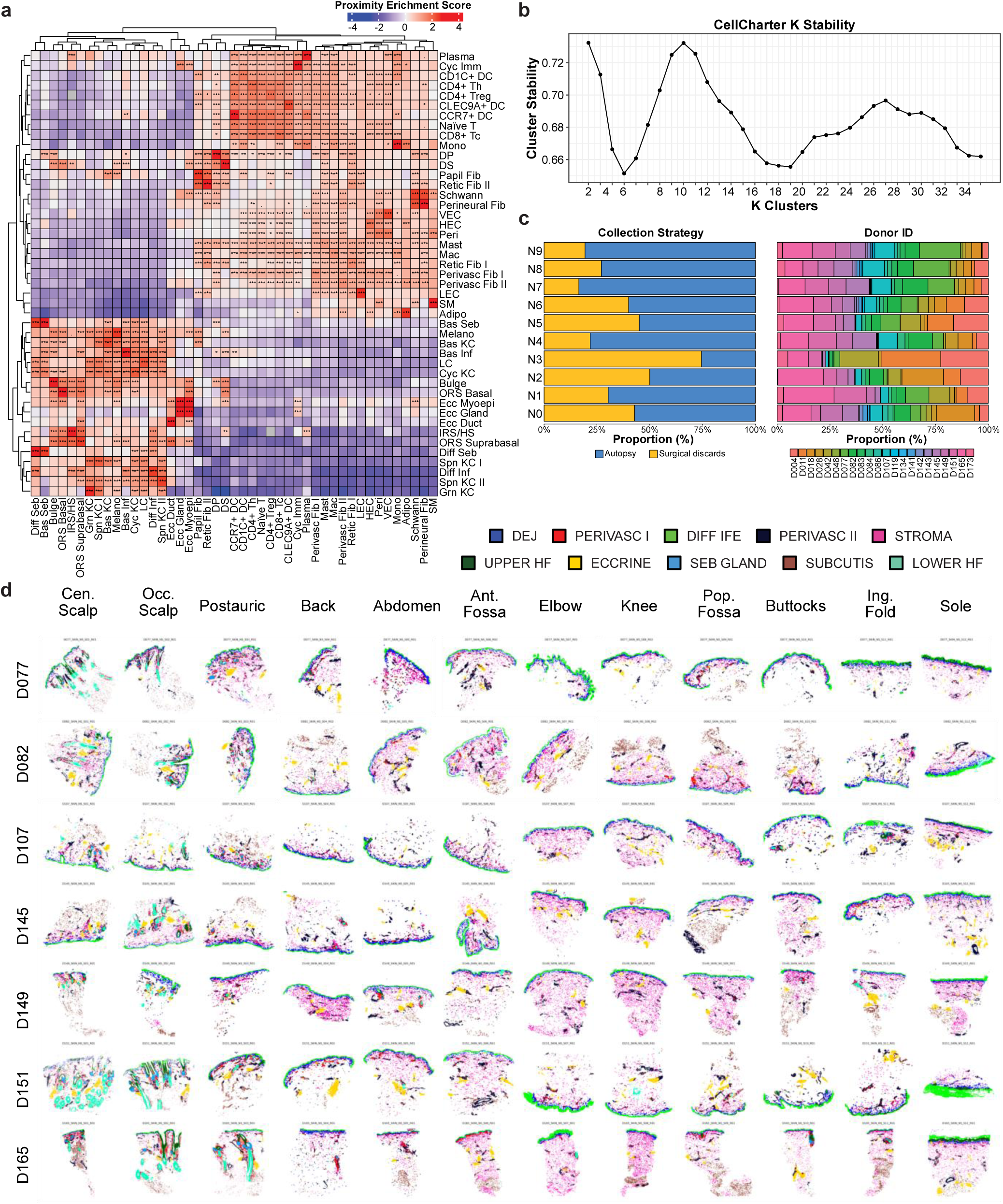
MERFISH identifies multicellular neighborhoods in human skin. **a.** Pairwise cell type proximity scores across all MERFISH samples. **b.** Cluster stability scores in reiterative clustering in CellCharter. Global maximum was attained at k = 10 clusters. **c.** Left, bar plots of each neighborhood’s sample composition between autopsy and surgical discards. Right, bar plots of each neighborhood’s sample composition by donor. **d.** All cells labeled by neighborhoods across MERFISH samples (n = 84 samples from 7 donors).

**Extended Data Figure 8.**
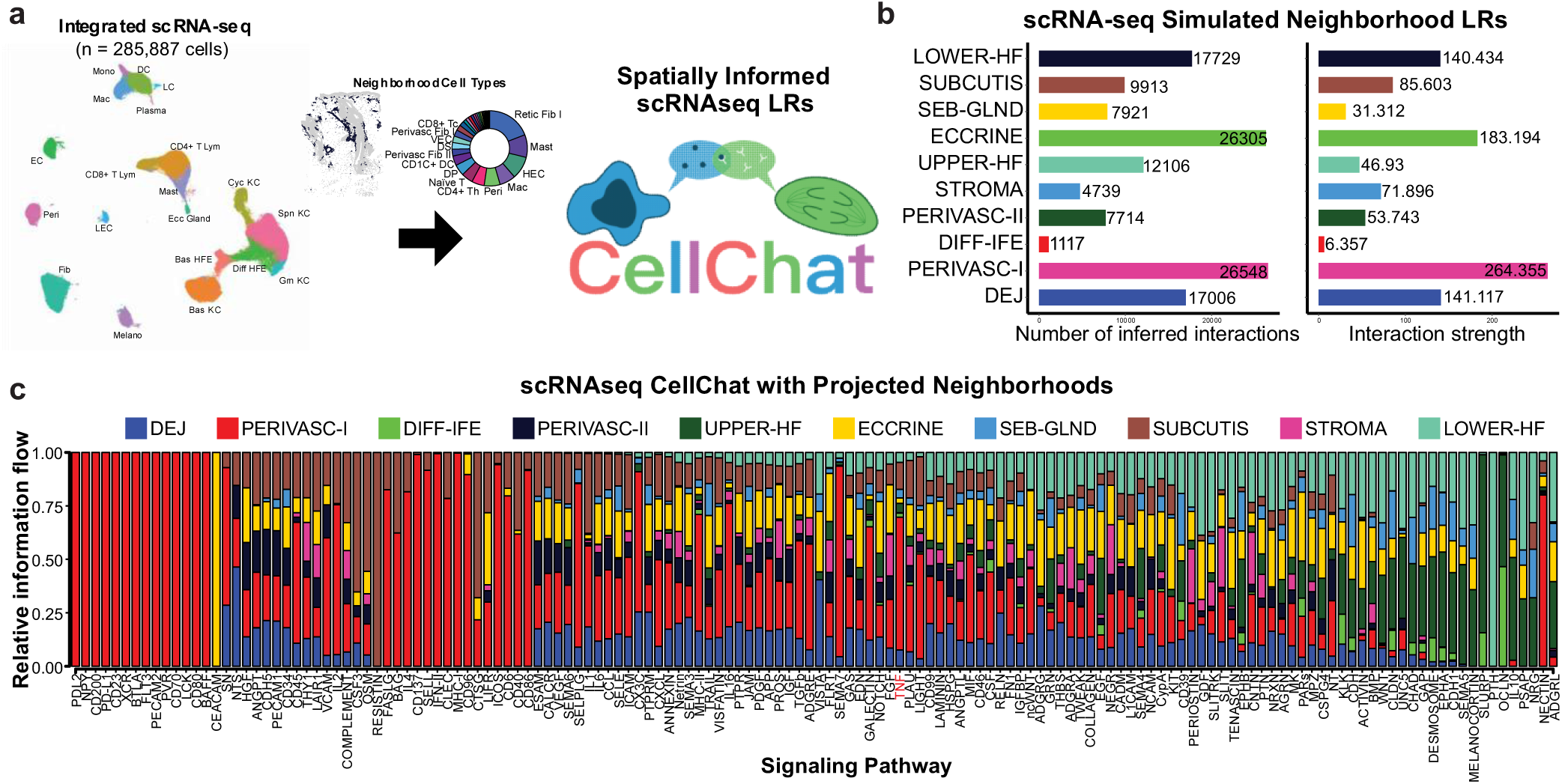
Spatially-informed ligand-receptor signaling across neighborhoods. **a.** Schematic of simulated scRNA-seq neighborhoods. **b.** Bar plots of inferred interactions (left) and interaction strength (right) of each neighborhood as determined by CellChat. **c.** Bar plots of neighborhood contributions to inferred active pathways. TNF is highlighted in red.

**Extended Data Figure 9.**
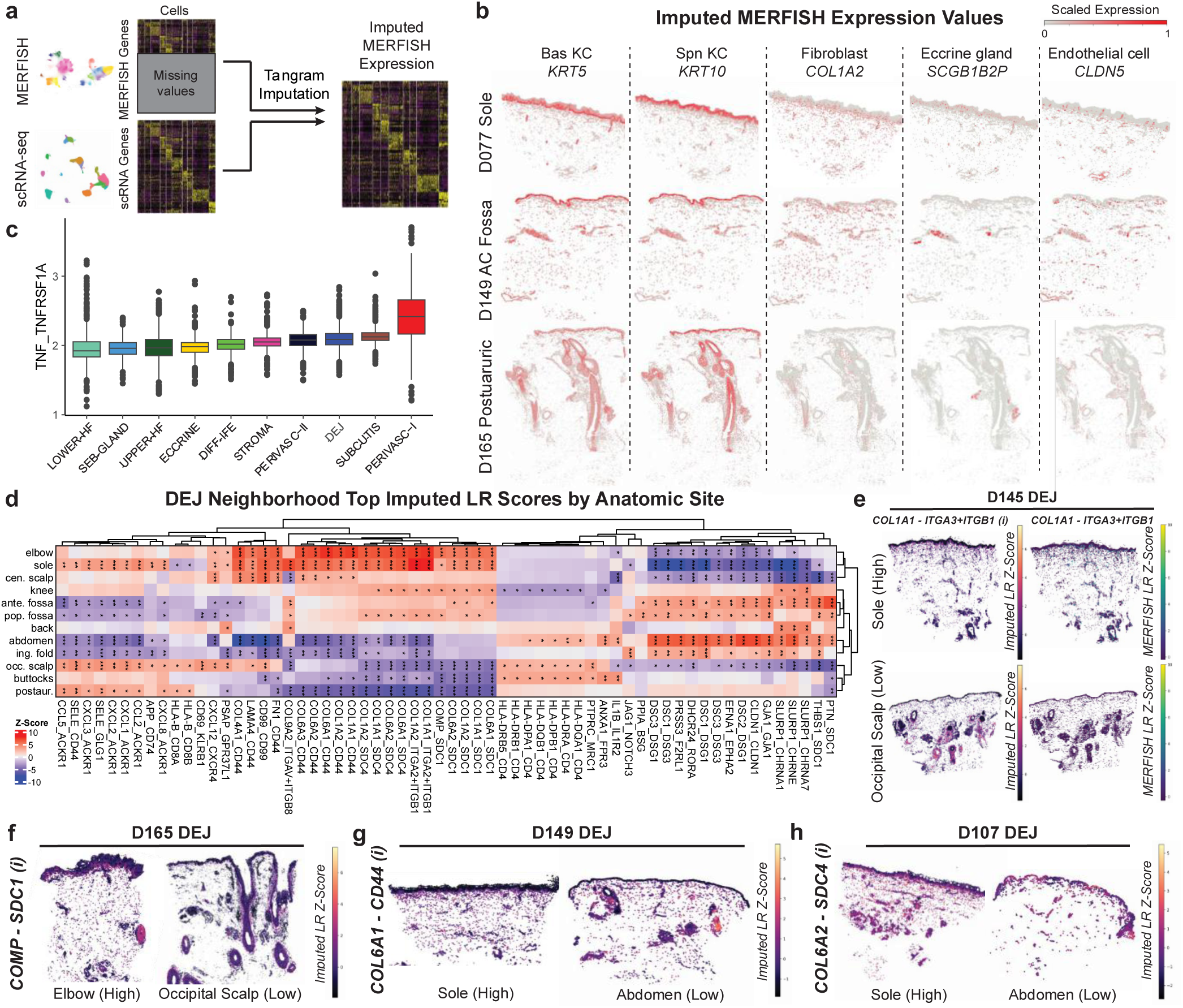
scRNA-seq imputation expands neighborhood neighborhood analyses transcriptome-wide. **a.** Schematic of imputation of MERFISH expression using scRNA-seq as a reference using Tangram. **b.** Imputed spatial expression of indicated marker genes from single cells in MERFISH data. **c.** Box plots of imputed TNF-TNFRSF1A LR scores per neighborhood across all samples. Boxes display median and top and bottom quartiles of the data. **d.** Heatmap of top imputed LR scores at each anatomic site in the dermo-epidermal junction (DEJ) neighborhood. *adjusted p < 0.05, **adjusted p < 0.01, ***adjusted p < 0.001, moderated t test. **e.** Imputed (i) and real MERFISH COL1A1-ITGA3+ITGB1 ligand-receptor (LR) scores at high (sole) and low (occipital scalp) scoring sites in D145. **f.** Imputed (i) COMP-SDC1 ligand-receptor (LR) scores at high (elbow) and low (occipital scalp) scoring sites in D165. **g.** Imputed (i) COL6A1-CD44 ligand-receptor (LR) scores at high (sole) and low (abdomen) scoring sites in D149. **h.** Imputed (i) COL6A2-SDC4 ligand-receptor (LR) scores at high (sole) and low (abdomen) scoring sites in D107.

**Extended Data Figure 10.**
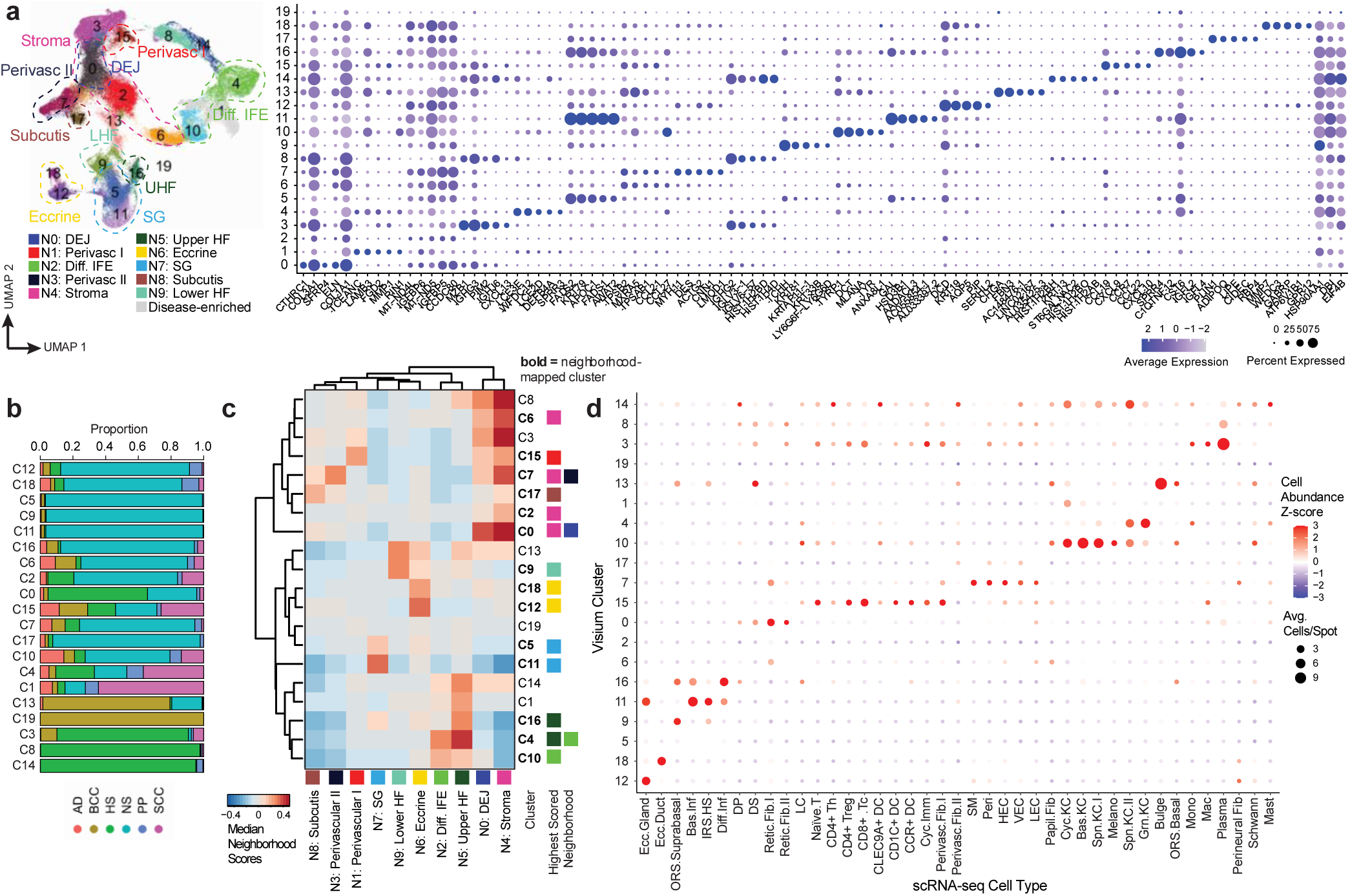
Mapping of MERFISH neighborhoods onto human skin diseases. **a.** Left, UMAP of all Visium spots (n = 142,515 spots) labeled by 20 unbiased clusters. Clusters are additionally annotated by MERFISH neighborhoods they resemble the most. Right, dot plot of top cluster markers. **b.** Bar plots of tissue state proportions comprising each unbiased spot cluster. **c.** Heatmap of median neighborhood scores (see Methods) for each cluster. Normal skin (NS) enriched clusters are bolded. **d.** Dot plot of cell2location deconvolustion of Visium spots, with cell type abundance and densities. Most clusters score for multiple cell types that correspond to cell types within MERFISH neighborhoods.

**Extended Data Figure 11.**
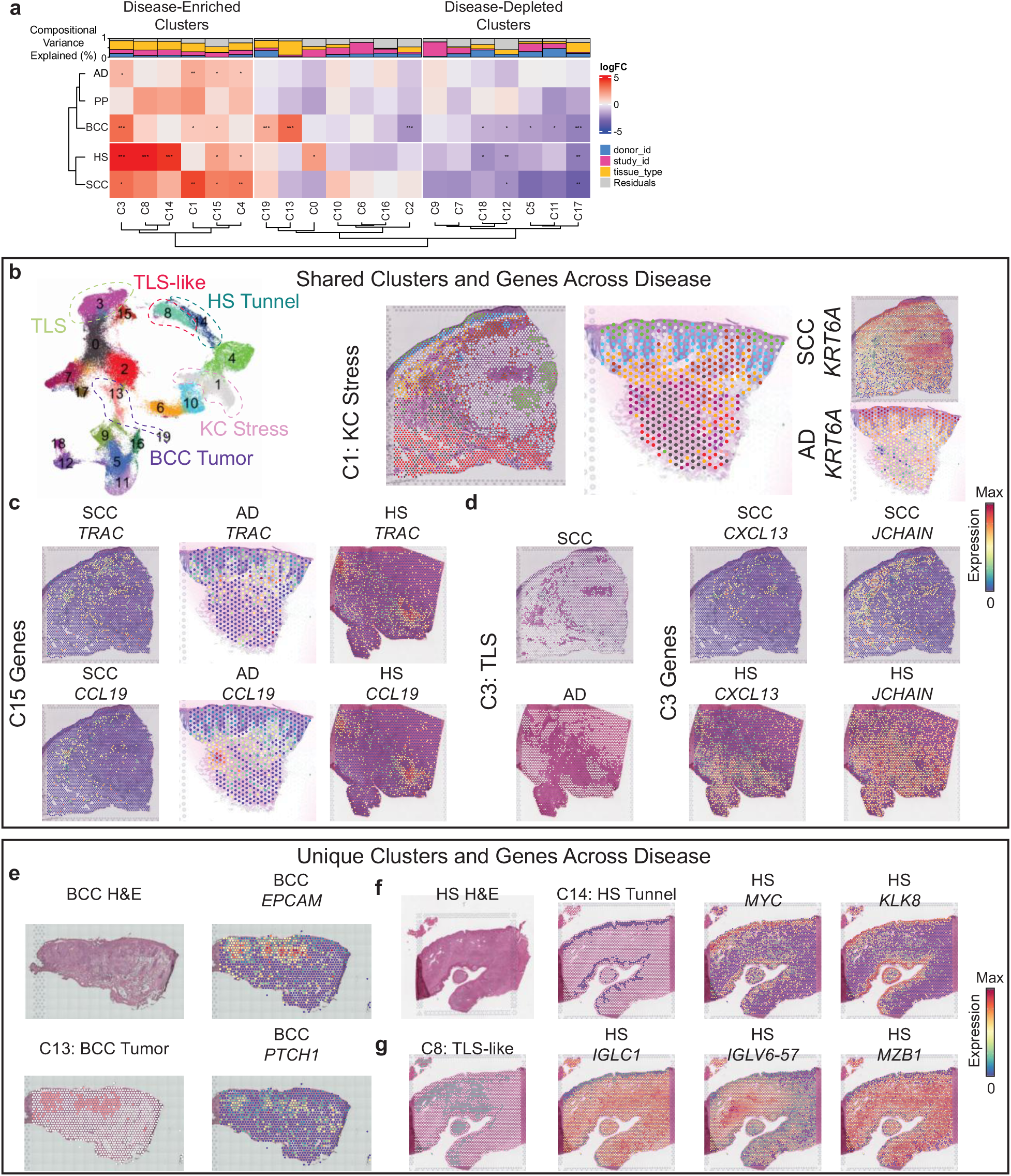
Perivascular immune-stromal remodeling in human skin disease. **a.** Heatmap of cluster abundances of diseased tissues relative to normal skin. *adjusted p < 0.05, **adjusted p < 0.01, ***adjusted p < 0.001, moderated t test. **b.** Left, UMAP of all Visium spots (n = 142,515 spots) labeled by 20 unbiased clusters and annotated for disease enriched clusters. Right, localization of C1 stressed KC cluster in SCC and AD samples with corresponding *KRT6A* marker expression. **c.** Localization of C15 perivascular marker gene expression in SCC, AD, and HS samples. **d.** Localization of C3 tertiary lymphoid structure (TLS) cluster in SCC and HS samples with corresponding *CXCL13* and *JCHAIN* marker expression. **e.** Localization of C13 BCC tumor cluster with corresponding *EPCAM* and *PTCH1* marker expression in BCC sample. **f.** Localization of C14 HS tunnel cluster with corresponding *MYC* and *KLK8* marker expression in HS sample. **g.** Localization of C8 TLS-like cluster with corresponding *IGLC1 IGLV6 57 an M B1* marker expression in HS sample.

## SUPPLEMENTAL TABLES

*Supplemental tables will be made available upon publication.*

**Table S1.** Human Skin Cohort Metadata.

**Table S2.** Human Skin MERFISH Panels.

**Table S3.** MERFISH Cluster Markers.

**Table S4.** scRNA-seq Cluster Markers.

**Table S5.** CellChat Results for MERFISH and scRNA-seq Neighborhoods.

**Table S6.** MERFISH and Tangram Ligand-Receptor Scoring.

**Table S7.** Visium Cluster Markers.

